# Immediate and deferred epigenomic signature of neuronal activation

**DOI:** 10.1101/534115

**Authors:** Jordi Fernandez-Albert, Michal Lipinski, María T. Lopez-Cascales, M. Jordan Rowley, Ana M. Martin-Gonzalez, Beatriz del Blanco, Victor G. Corces, Angel Barco

## Abstract

Activity-driven transcription plays an important role in many brain processes, including those underlying memory and epilepsy. Here, we combine the genetic tagging of neuronal nuclei and ribosomes with various sequencing-based techniques to investigate the transcriptional and chromatin changes occurring at hippocampal excitatory neurons upon synchronous activation during status epilepticus and sparse activation during novel context exploration. The transcriptional burst, which affects both nucleus-resident non-coding RNAs and numerous protein-coding genes involved in neuroplasticity, is associated with a dramatic increase in chromatin accessibility of activity-regulated genes and enhancers, *de novo* binding of activity-regulated transcription factors, augmented promoter-enhancer interactions, and the formation of gene loops that bring together the TSS and TTS of strongly induced genes to sustain the fast re-loading of RNAPII complexes. Remarkably, some chromatin occupancy changes and interactions remain long after neuronal activation and may underlie the changes in neuronal responsiveness and circuit connectivity observed in these neuroplasticity paradigms.

## Introduction

Activity-dependent transcription is a key part of the neuronal response to synaptic stimulation and is essential for the transition from short-to long-term forms of neuronal plasticity ^1, 2^. As a result, its deregulation is an important feature of many neurological diseases associated with cognitive dysfunction ^3^. This is the case for epilepsy, a severe neurological condition that predisposes the patient to recurrent unprovoked seizures or transient disruption of brain function due to abnormal and uncontrolled neuronal activity, and can lead to brain damage, disability and even death ^4^. Studies in experimental models of epilepsy have shown that a single epileptic seizure episode – *aka* status epilepticus (SE) – evokes a strong transcriptional response in the hippocampus that includes the transient transcription of immediate-early genes (IEGs) ^5–7^, rapid changes in chromatin marks and nuclear structure ^8–12^, and a delayed transcriptional response affecting many synaptic effector genes ^1, 13^. Beyond this pathological setting, activity-dependent transcription is also thought to contribute to the changes in circuit connectivity and neuronal responsiveness that are necessary for memory formation ^14, 15^. Thus, it has been hypothesized that the formation of long-term memory, which requires the activation of tightly regulated transcriptional programs during learning, relies not only on synaptic changes but also on modifications in the chromatin of activated neurons ^16, 17^. These epigenetic changes may contribute to long-lasting or permanent changes in the expression and responsiveness of genes involved in synaptic function, thereby representing a sort of genomic memory.

New sequencing techniques have unveiled novel processes that contribute to the complex transcriptional response to neuronal activation, such as activity-regulated intron skipping, double-strand breaks, and the production of different species of non-coding RNAs (ncRNAs), including enhancer (eRNAs) and extracoding RNAs (ecRNAs) ^18–24^. Despite this progress, an integrated view of the transcriptional and epigenomic changes triggered by neuronal activation *in vivo* is still lacking, partially because of the special challenges derived from brain complexity and cellular heterogeneity ^1^. For instance, we do not know whether the processes mentioned above, most of which have only been described *in vitro*, also occur in the brain of behaving animals. We also still ignore the mechanisms that sustain the activity-driven transcriptional burst and the nature of the afore predicted genomic memory.

Here, we introduce methods to efficiently profile the transcriptome and epigenome of defined neuronal populations in the adult brain. Using these methods we were able to generate the first high-resolution, multi-dimensional and cell-type-specific draft of transcriptional and chromatin alterations associated with neuronal activation *in vivo*, both under physiological and pathological conditions. Our genome-wide screens and analyses provide mechanistic insight into both the rapid changes in gene expression that are initiated after neuronal activity and the more enduring changes that may underlie processes such as epilepsy and memory formation.

## Results

### Nuclear tagging enables neuronal-specific genomic screens and reveals broad changes to the nuclear transcriptome upon SE

To investigate the transcriptional and chromatin changes specifically occurring in forebrain principal neurons upon activation, we produced mice in which the nuclear envelope of these neurons is fluorescently tagged (Fig. 1A-B). This genetic strategy combined with fluorescence activated nuclear sorting (FANS) allowed us to isolate this neuronal nuclei type from adult brain tissue (**Fig. S1A-C**). After FANS, sorted nuclei can be used in different applications, from nuclear RNA (nuRNA) quantification (**Fig. S1D**) to chromatin immunoprecipitation (ChIP) (**Fig. S1E**) and enzymatic treatments like the assay for transposase-accessible chromatin (ATAC) (**Fig. S1F**). FANS represents a substantial improvement over cell sorting because it prevents the stress response triggered by neuronal dissociation that can occlude activity-induced changes. Moreover, since our method does not require the use of antibodies against GFP, it can be combined with immunolabeling for secondary sorting to increase the specificity of the cellular population under investigation.

**Figure 1.**
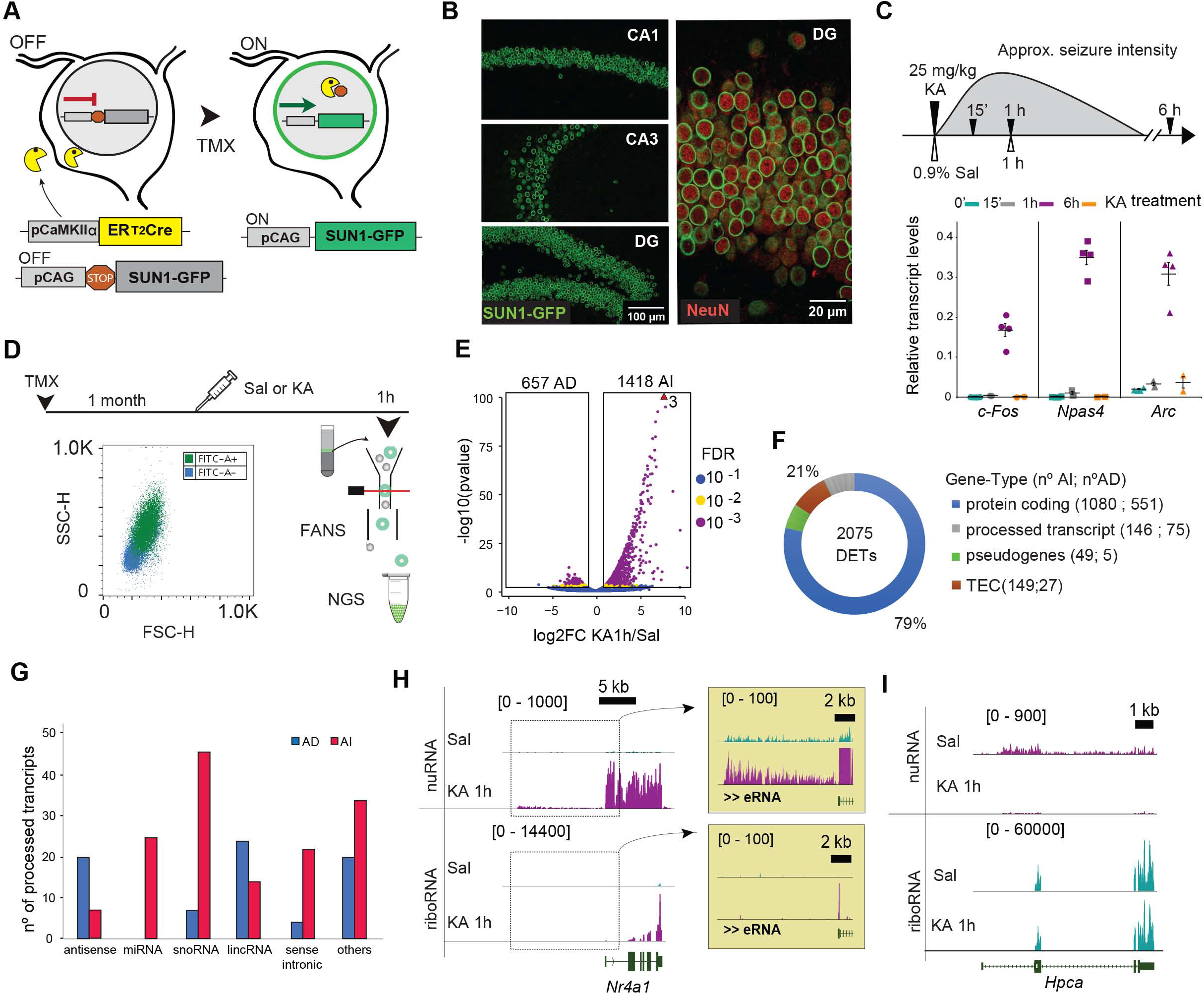
SE triggers broad changes of the nuclear transcriptome. **A**. Crossing of the TMX-inducible Cre-driver line CaMKII-creERT2 with mice that express SUN1-GFP in a Cre-recombination-dependent manner allows the tagging of the nuclear envelope in forebrain principal neurons for subsequent FANS-based isolation. **B**. Confocal images of principal hippocampal neurons in the CA1, CA3 and DG subfields stained against GFP (green). **C**. Top: Scheme depicting seizure strength and duration in KA-treated animals. Mice were injected with KA (25 mg/kg) and sacrificed at various time points. As a reference, control mice were treated with saline and sacrificed 1 h after treatment. Bottom: RT-qPCR assays show transient IEG induction. **D**. Experimental design for KA-induced neuronal activity, nuclei isolation and application of next generation sequencing (NGS) techniques. **E**. Volcano plot showing the significance and p-value distribution after differential gene expression analysis. AD: Activity Depleted; AI: Activity Induced. **F**. Gene biotype classification for DETs 1 h after KA in the nuRNA-seq (left) and riboRNA-seq (right) screens. TEC: To be Experimentally Confirmed. **G**. Number of processed transcript species detected for DETs 1 h after KA in the nuRNA-seq screens. miRNA: micro RNA; snoRNA: small nucleolar RNA; lincRNA: long interspersed non-coding RNA. **H**. NuRNAseq detects activity-induced eRNA and other nucleus-resident ncRNAs. We also present the riboRNAseq tracks for comparison. Values represent counts in RPM (read per million). **I**. Genomic track for the gene encoding hippocalcin (*Hpca*), a calcium-binding protein that is highly expressed in the hippocampus and downregulated upon SE. Values represent counts in RPM. See also Figure S1.

Exposure to the glutamate receptor agonist kainic acid (KA) is often used to model SE in experimental animals. Nuclear envelope-tagged mice were treated with 25 mg/Kg of KA, which causes a strong and synchronous activation in all the hippocampal subfields leading to a robust but transient induction of IEGs (Fig. 1C). We used FANS to isolate nuRNA from hippocampal principal neurons of KA- and saline-treated mice (Fig. 1D and **S1G**). This screen detected over 2,075 differentially expressed transcripts (DETs) (Fig. 1E and **Table S1**), unveiling a broader impact and less skewed distribution than previous studies of activity-induced transcription ^2^. We observed the induction of a rich variety of transcripts, including eRNAs and other nucleus-resident species (Fig. 1F-H). These profiles did not present the typical enrichment for 3’ sequences observed in mRNA-seq and covered both exons and introns (Figs. 1H **and S1H**). These results indicate that nuRNA-seq is particularly well suited for studying dynamic transcriptional responses because the signal precisely reflects the time point in which the transcripts are produced. As a result, nuRNA-seq provides a finer detection of activity-induced changes, particularly for transcripts that are already expressed in the basal state. It also unveils the transient downregulation of hundreds of transcripts (Fig. 1E, I), including many genes involved in basal metabolism (**Fig. S1I**), which suggests that the activity-induced transcriptional burst hijacks the transcriptional machinery leading to a transient shutdown of basal transcription.

### Compartment-specific transcriptomics reveals the uncoupling of the SE-induced neuronal transcriptome and translatome

We next used the same genetic approach to tag ribosomes and identify the pool of mRNAs that are translated in hippocampal excitatory neurons in response to seizures (i.e., the activity-induced neuronal translatome) (Figs. 2A-C and **S2A**). Translating ribosomal affinity purification (TRAP) ^25^ led to an enrichment for neuron-specific transcripts and depletion of glia-specific transcripts (Figs. 2D and S2B). The sequencing (referred to as riboRNA-seq) and analysis of TRAPped mRNAs retrieved 189 differentially translated genes (DTGs) (Figs. 2E**, S2B-E** and **Table S2**). Most of these DTGs are protein-coding (Figs. 2F and **S2F**) and relate to Gene Ontology (GO) functions such as *Transcription cofactor activity, Kinase activity, Histone acetyltransferase* and *Chromatin binding*, being the nucleus the most enriched cellular component (**Fig. S2G**, red bars). Notably, the small set of activity-depleted mRNAs is also related to transcriptional regulation (**Fig. S2G**, blue bars), which further underscores the importance of transcriptional and epigenetic regulation in activity-dependent processes.

**Figure 2.**
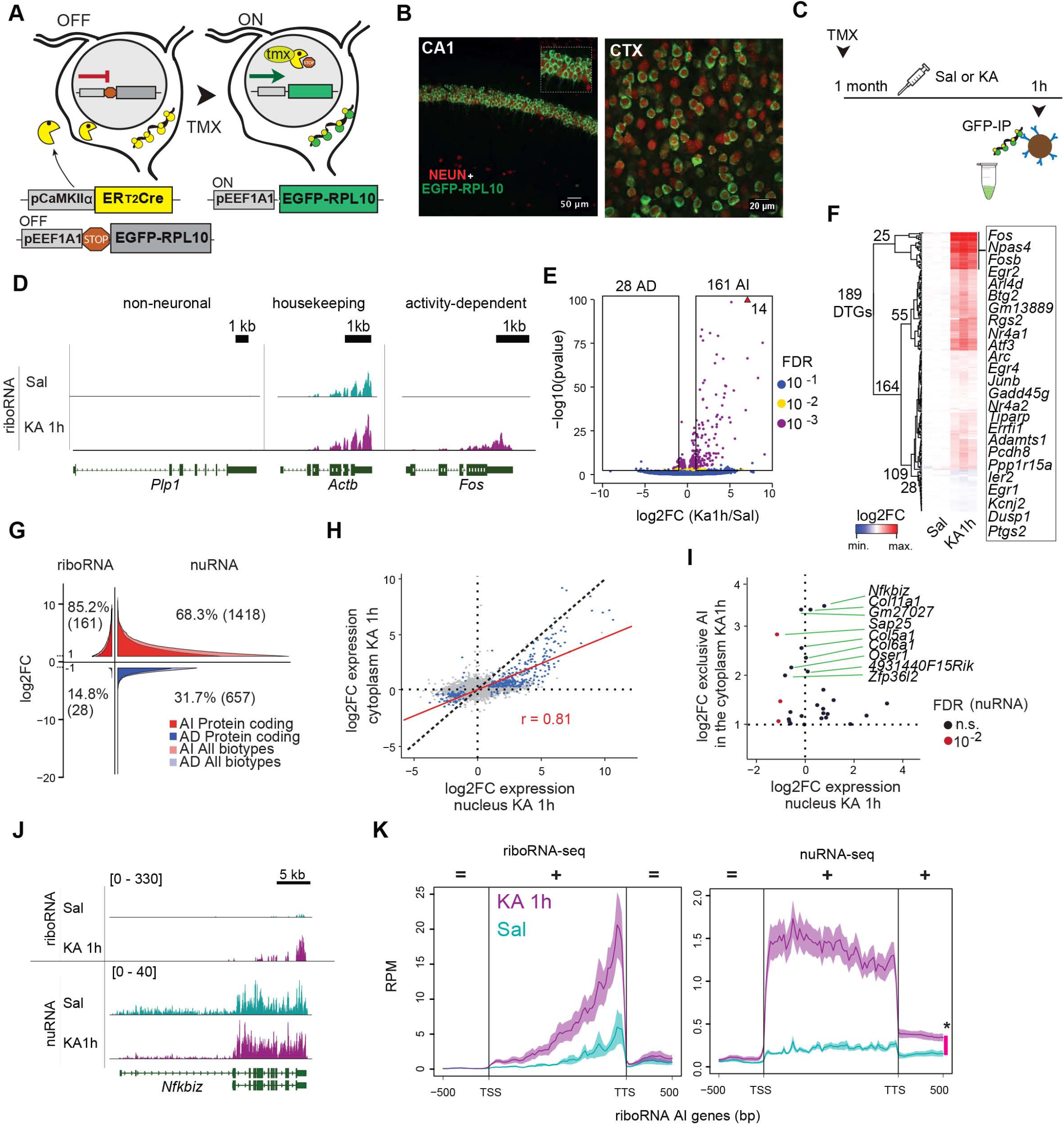
Compartment-specific transcriptomics reveals changes in neuronal transcripts production and translation. **A**. Crossing the TMX-inducible Cre-driver line CaMKII-creERT2 with mice that express GFP-L10a in a Cre-recombination-dependent manner allows the tagging of ribosomes in forebrain principal neurons for subsequent IP-based isolation. **B**. Confocal images of principal neurons in the CA1 subfield and cortex stained against GFP (green) and the neuron marker NeuN (red). **C**. Experimental design for KA-induced neuronal activation and ribosomal RNA isolation. **D**. Genomic snapshots for representative examples of glia-specific (*Plp1*), housekeeping (*ActB*) and activity-induced (*Fos*) genes. Values indicate the levels of counts in RPM. **E**. Volcano plot showing the significance and p-value distribution after differential transcript abundance analysis. AD: Activity Depleted; AI: Activity Induced. **F**. Heatmap of fold changes in DTGs retrieved in the riboRNA-seq screen. **G**. Density distribution of DTGs and DETs in the riboRNAseq and nuRNAseq screens, respectively. Colors indicate the distribution of protein coding *vs* all biotypes. **H**. Correlation of fold change (FC) values for genes significantly regulated (FDR < 0.1, 372 genes) in both datasets (blue). All other detected genes are labeled in gray. **I**. Scatter plot of log2 FC for genes exclusively upregulated in the cytoplasm (FDR < 0.1) and its FC value in the nucleus (colors indicate FDR significance in nuRNA-seq analysis; n.s.: FDR > 0.1 in nuRNA-seq). **J**. Comparison of nuRNA-seq and riboRNA-seq tracks in *Nfkbiz*. The vertical scale shows counts in RPM. **K**. Metagene mapability profiles for riboRNA-seq and nuRNA-seq samples at genes upregulated in both datasets. The red bar to the right indicates the significant difference downstream of the TTS in nuRNA-seq. See also Figure S2.

The availability of neuronal-specific riboRNA-seq and nuRNA-seq data provides a unique opportunity for the dissection of RNA-related mechanisms involved in the response to activation. First, the comparison of the number, magnitude and distribution of the changes confirmed the special suitability of nuRNA-seq for transcriptional dynamics studies (Figs. 1G-H and 2G). Second, despite these differences, we observed a robust correlation between changes in both screens demonstrating the coordinated regulation of RNA transcription and export upon neuronal activation (Fig. 2H). Third, we identified a small number of transcripts that escape the general correlation. These mRNAs show an increase in riboRNA-seq profiles but no change, or even a decrease in nuRNA-seq profiles. This pattern suggests that activity-dependent regulation of these transcripts occurs at the translational or RNA stability levels rather than at transcriptional level (Fig. 2I). Notably, some of these genes are known to be post-transcriptionally regulated in other cell types ^26^ (Figs. 2J and **S2H**, discordant). This is the case with *Nfkbiz*, which encodes an atypical member of the NFκB family and harbors a translational silencing element (TSE) in the 3’ UTR that destabilizes the mRNA ^27, 28^ Fourth, we observed striking differences in the transcription termination site (TTS) of IEG transcripts obtained after nuclear and ribosome-bound RNA isolation. The nuclear transcripts were consistently longer, extending several hundred base pairs after the annotated TTS (Fig. 2K, red bar). This observation indicates that activity-induced ecRNAs, which have been postulated to prevent gene inactivation in embryonic neuronal cultures ^19^, are also produced in the adult brain. Overall, these findings underscore the value of parallel compartment-specific transcriptome analyses to unveil novel regulatory mechanisms.

### Transcriptional bursting causes a dramatic increase in accessibility at IEGs

To precisely correlate transcriptional differences with changes in chromatin occupation and TF binding, we next assessed chromatin accessibility by ATAC-seq ^29^. The innovative combination of FANS and ATAC-seq analyses revealed that the chromatin accessibility profile of hippocampal neurons (**Fig. S3A-C**) is dramatically altered during SE (Fig. 3A). The differential accessibility (DA) screen retrieved more than 30,000 differentially accessible regions (DARs) (Fig. 3B and **Table S3A**), providing a deeper survey of activity-dependent changes than previous analyses ^30^ (**Fig. S3D-F**). While KA-induced chromatin accessibility was primarily observed within gene regions, particularly within close proximity of the TSSs (< 1 Kb), chromatin closing occurred more at intergenic regions (Fig. 3C). We used binding and expression target analysis (BETA) to integrate ATAC-seq and RNA-seq data ^31^ and found that the increase in chromatin accessibility throughout the gene body is an excellent predictor of transcriptional activation (p < 3.86×10^−36^, Fig. 3D). ATAC-seq data correlated with changes in the riboRNA-seq and nuRNA-seq screens (**Fig. S3G**), but correlation was stronger and depended on more genes in the case of nuRNA-seq (**Fig. S3H** and **Table S3B**). These comparisons revealed a very prominent increase in accessibility at the gene body of activity-induced genes that extended beyond the TTS and narrowly matched transcript production (Fig. 3E).

**Figure 3.**
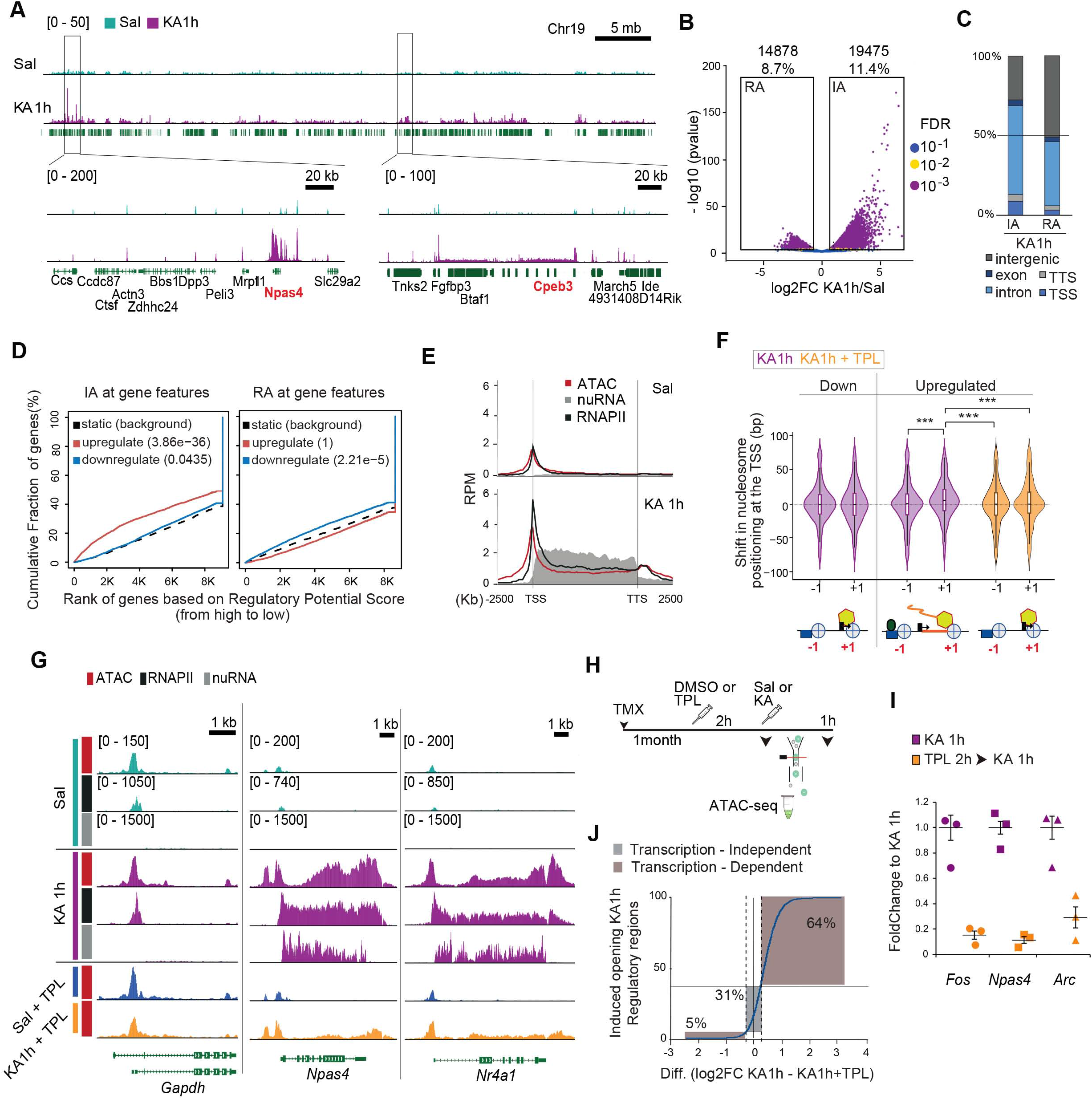
Neuronal activation causes a dramatic increase in accessibility at activity-regulated genes associated with the transcriptional burst. **A**. Genomic profile at the chr19 locus containing the activity-induced genes *Npas4 and Cpeb3* (in red). **B**. Volcano plot showing the significance value distribution after DAR analysis; upper values indicate number of regions and percentage of all the accessible regions detected. RA: Reduced Accessibility; IA: Increased Accessibility. **C**. Distribution of genomic features along differentially accessible regions. **D**. Predictive analysis of activating/repressive function for accessibility changes at regions displaying increased (IA) or reduced accessibility (RA) within the gene body and correlation with changes in nuclear transcript levels. **E**. Metagene plot for ATAC-seq, RNAPII-ChIP-seq and nuRNA-seq signals at upregulated (log2FC > 3, FDR < 0.1) genes 1 h after saline or KA administration. **F**. Violin plot showing the shift in the distribution of nucleosome positioning at the TSS of downregulated and upregulated genes 1 h after KA (purple) or after KA+TPL treatment (orange). ***: p-adjusted < 0.00001 in *pos hoc* Dunn test for Kluskal-Wallis with multiple comparisons. **G**. Top: Genomic snapshots of ATAC-seq, nuRNA-seq and RNAPII-ChIP-seq profiles in control and activity-regulated genes 1 h after saline or KA administration. Bottom: Impact of TPL in ATAC-seq profiles. Values indicate the levels of counts in RPM (read per million). **H**. Experimental design for the TPL blockage of KA-induced changes. **I**. RT-qPCR assays show the inhibition of IEG induction by TPL after KA-induced neuronal activation. **J**. Distribution of the difference of FC for “KA-1h” and “TPL+KA-1h” at DARs KA-1h. The sites considered to be transcription independent (log2 FC difference larger than ±0.25) are in grey, while the ones considered to be transcription dependent are in green. See also Figure S3.

We next investigated the relationship between changes in accessibility, transcription and RNA polymerase II complex (RNAPII) binding. Experiments in neuronal cultures have suggested that IEGs may have pre-assembled RNAPII at the TSS in the basal state ^32^. RNAPII poising is however not evident *in vivo*. Although we detected a modest presence of RNAPII at the promoter of rapid response genes in saline-treated mice, SE triggered a robust *de novo* binding (Fig. 3E). Nucleosome positioning at the TSS of activity-regulated genes showed a shift of the +1 nucleosome in activity-induced genes, also reflecting the *de novo* entrance of RNAPII (Fig. 3F) ^33^. In addition to promoter binding, neuronal activation increased RNAPII occupancy of the gene body and the TTS in activity-regulated genes matching the production of extended nuclear transcripts (Fig. 3E, G) and enhanced intragenic accessibility (**Fig. S3I**). These results suggest that the enhanced accessibility at IEGs reflects the continuous passage of the RNAPII. To assess this hypothesis, we examined the effect of triptolide (TPL), a drug that inhibits transcription initiation ^34, 35^, in chromatin accessibility changes (Fig. 3H). Both IEG induction (Fig. 3I) and increased accessibility in the gene body of upregulated genes (Fig. 3G and **Table S3C**) were reduced in animals treated with TPL before KA administration. The impact of TPL on activity-dependent DA was more prominent in the genes that show the largest changes (**Fig. S3J**) and stronger in gene bodies than TSSs (**Fig. S3K**), thereby confirming that most of these changes are directly derived from the traffic of RNAPII during the transcriptional burst (**Fig. S3L**). Administration of TPL also prevented the nucleosome shift at the TSS caused by the entrance of new RNAPII complexes (Fig. 3F). Interestingly, our analyses show that about 30% of the regions with altered accessibility in response to KA were not affected by TPL (Fig. 3J), indicating that other mechanisms, such as the *de novo* binding of activity-regulated transcription factors (TF) and the recruitment of co-activators, also contribute to activity-dependent changes in the chromatin.

### TF and transcriptional co-activator binding at activity-regulated enhancers

Similar to TSSs and intragenic sequences (Fig. 3D), SE-induced changes at extragenic regions (> 1 Kb from gene body) also correlates with transcriptional changes in the proximal genes (**Fig. S4A**). To explore in greater detail these changes, we classified the accessible extragenic regions in the chromatin of mature excitatory neurons into promoters and putative enhancers distinguishing between regions with a stronger or weaker epigenetic signature (Fig. 4A). Promoters were characterized by the prominent presence of RNAPII and H3K4me3, along with a depletion of H3K4me1. In contrast, putative enhancers presented enrichments for the histone post-translational modifications (HPTM) H3K4me1 and H3K27ac and binding of the transcriptional co-activator and lysine acetyltransferase CREB-binding protein (CBP). About 10% of these promoter regions and 25% of the putative enhancer regions showed changes in accessibility during SE (**Fig. S4B**). Notably, these changes were consistent with ChIP-seq profiles for the binding of RNAPII and CBP upon SE, indicating that the changes are driven by the activity-dependent recruitment of transcriptional complexes. While promoters and strong enhancers showed robust activity-dependent binding of both RNAPII and CBP, weak enhancers only showed CBP binding (**Fig. S4C**), indicating that they may be poised for activation ^36^. Consistent with this view, we detected an activity-dependent increase in H3K27 acetylation at the same sites (**Fig. S4D**) in primary cultures of embryonic cortical neurons stimulated with KCl ^37^. Furthermore, activity-dependent CBP recruitment and increased H3K27 acetylation occurred at both TPL-sensitive and TPL-resistant sites (**Figs. S4E-F**), which suggests that CBP recruitment and enhancer activation may be independent of RNAPII action. Intriguingly, 75% of the activity-induced (AI)-transcripts are associated with activity-driven changes in accessibility (Fig. 4B-C), but only 25% of promoters and 10% of enhancers displaying increased accessibility are linked to AI genes (**Fig. S4G**). This discrepancy indicates that synaptic activity triggers a large number of chromatin changes that are not directly associated with transcription in the nearest gene.

**Figure 4.**
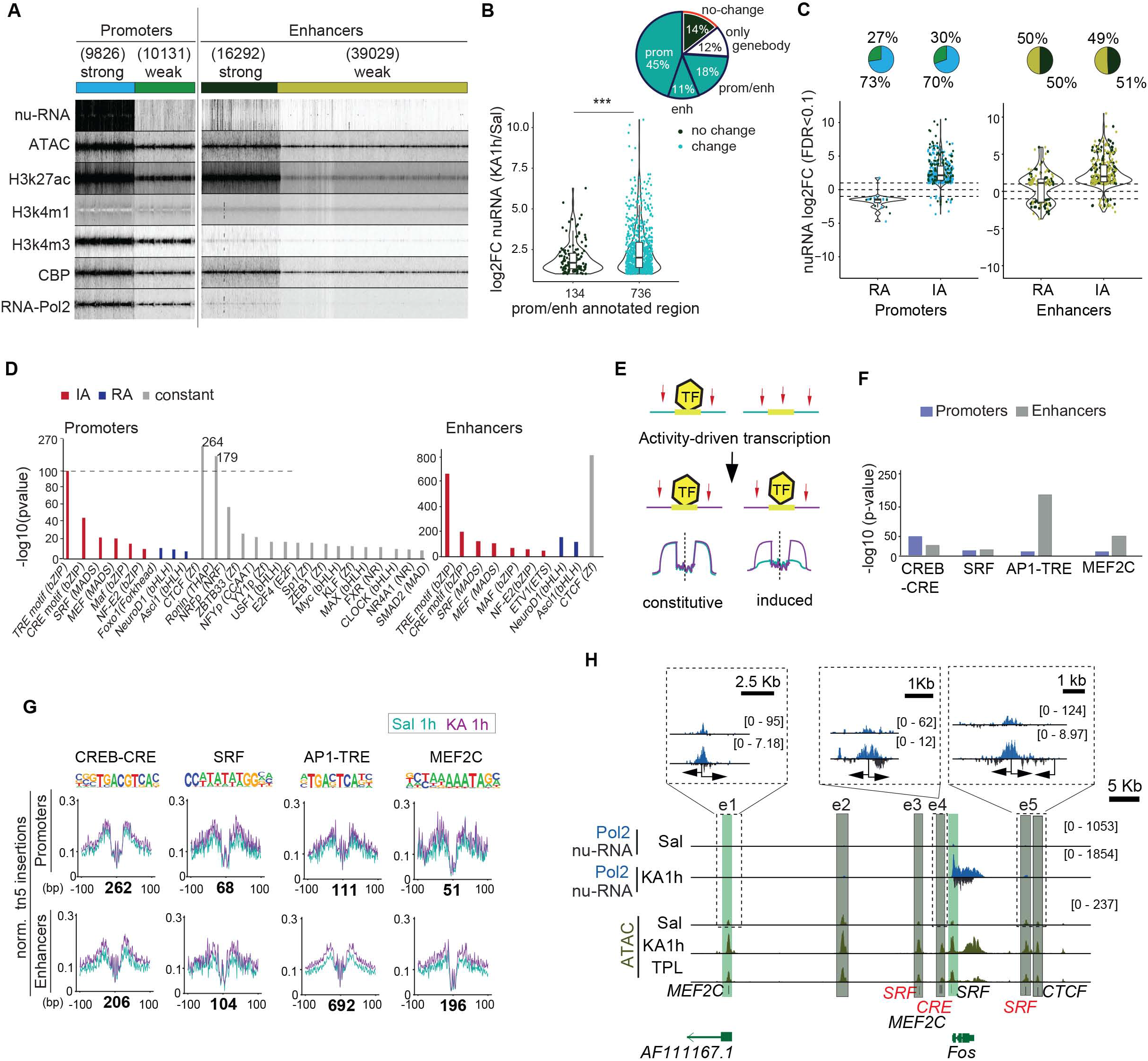
Neuronal activation induces TF-binding at extragenic sites. **A**. K-mean clustering for accessible extragenic regions (± 5 Kb) generated using the profiles of ATAC-seq, nuRNAseq, and ChIP-seq for RNAPII, CBP, H3K27ac, H3K4m3 and H3K4m1 in hippocampal chromatin of naïve animals (this study and ^51, 52^). **B**. Violin plot shows the log2FC distribution and number of upregulated nuclear transcripts with promoter/enhancer annotations that change (cyan) or do not change (black) during SE. ***: p-adjusted value = 2.10^−16^ in Kruskal-Wallis test. The sector graph indicates the percentage of regions according to their annotation and change upon SE. **C**. Violin plots show the log2FC distribution of activity-regulated genes in KA-1h annotated with reduced (RA) or increased accessibility (IA) sites at promoters and enhancers. Note the detection of opposing changes in chromatin accessibility and transcript levels suggests the activity-dependent release of transcriptional repressors. Sector plot shows the percentage of strong and weak enhancers annotated at those genes. **D**. Motif enrichment comparison at detected promoter/enhancer accessible regions from opening (blue), closing (red) and constant (gray) regions. **E**. Scheme of digital footprinting analysis. Red arrows indicate sites accessible to tn5 cutting. Occupied TF-binding sites are more protected than the immediate surrounding. **F**. Motif enrichment significance at detected footprints in IA-promoters and -enhancers. **G**. Digital footprinting at enriched motifs and number of motifs detected (values correspond to normalized tn5 insertions). **H**. As an example, the snapshot shows ATAC-seq and RNAPII binding at regions flanking *Fos*. in saline, KA-1h and TPL+KA-1h samples (values in RPM). Footprints are labeled (red: lower stringency). The insets zoom on enhancers with detected eRNAs.

We next investigated the abundance of TF binding sites (TFBS) at accessible regions that show a significant increase (IA), reduction (RA), or no change in accessibility (constant) upon SE. IA promoters and enhancers were enriched for motifs recognized by TFs involved in neuronal plasticity processes such as AP1 (TRE) and CREB (CRE) ^2^; RA promoters and enhancers, on the other hand, showed enrichment for motifs associated with neuronal identity such as NeuroD1 (Fig. 4D). This result pinpoints again to a competition between activity-regulated genes and other highly expressed neuronal genes for the basal transcriptional machinery. Interestingly, constant promoters and enhancers were both enriched in CTCF binding sites, suggesting that the general architecture of the chromatin in neurons is not severely altered upon activation. To directly assess the occupancy of these sites before and during SE, we analyzed their digital footprints in ATAC-seq profiles (Fig. 4E). Occupancy by the activity-regulated TFs CREB, SRF, AP1 and MEF2C was detected in promoter and enhancer regions. While the footprint for the CREB family was stronger in promoters than in enhancers, the footprints of AP1 and MEF2C presented the opposite pattern (Fig. 4F-G). Upon SE, constitutive activity-regulated TFs, such as SRF and CREB, did not show large changes in their footprint although we could detect some *de novo* binding at enhancer regions (Figs. 4G-H and **S4H**). In contrast, AP1 showed a robust increase in occupancy at enhancer regions (Figs. 4F-G, and **S4H**) that, consistent with the *de novo* synthesis of AP1 proteins, was highly sensitive to TPL (**Fig. S4I-J**). ChIP-seq data for CREB, SRF and Fos in cortical neurons demonstrate the occupancy of these sites by the corresponding TFs after KCl stimulation ^18, 37^, suggesting that both modes of activation trigger the same genomic mechanisms (**Fig. S4K**). Consistent with the recently postulated role of the acquisition of signal-regulated DNA elements in gene regulatory regions in the evolution of cognitive abilities ^38^, these footprints are evolutionary conserved (**Fig. S4L**).

### Physiological and pathological neuronal activation share common mechanisms but trigger distinct epigenomic signatures

To investigate whether the genomic events described above are exclusive to pathological neuronal over-activation or can also be observed after neuronal activation in physiological conditions, we next explored chromatin changes triggered by the exploration of a novel and rich spatial context. This experience is known to induce IEG transcription in sparse neuronal assemblies in the hippocampus of rodents ^39^ (Fig. 5A). Nuclear envelope-tagged mice were subjected to novelty exploration (NE) for 1 h; we next used FANS and Fos immunostaining to isolate the small percentage of neuronal nuclei responding to this experience (25,000 Sun1^+^/Fos^+^ nuclei from the hippocampi of 6 novelty-exposed mice, Figs. 5B and **S5A**). We also isolated the same number of Sun1^+^/Fos^−^ nuclei for direct comparison of activated and non-activated neurons from the same animal. Both types of nuclei were processed for ATAC-seq to produce the first map of learning-related chromatin accessibility changes. Although the comparison of Fos^+^ and Fos^−^ neurons revealed less numerous and more modest changes (Fig. 5C**, S5B** and **Table S4**), there was a large overlap between the NE- and SE-induced DARs. Principal component analysis (PCA) showed that Fos^+^ nuclei clustered with the KA-1h samples, while Fos^−^ nuclei clustered with the samples of saline-treated mice (**Fig. S5C-D**). Similar to our SE study, chromatin accessibility changes in gene bodies correlated with nuclear transcript induction upon NE ^40^ (**Fig. S5E**), and digital footprinting revealed a clear enrichment for AP1 and Mef2c binding at activity-regulated enhancers, albeit less TF motifs were detected in NE than in SE (Fig. 5D).

**Figure 5.**
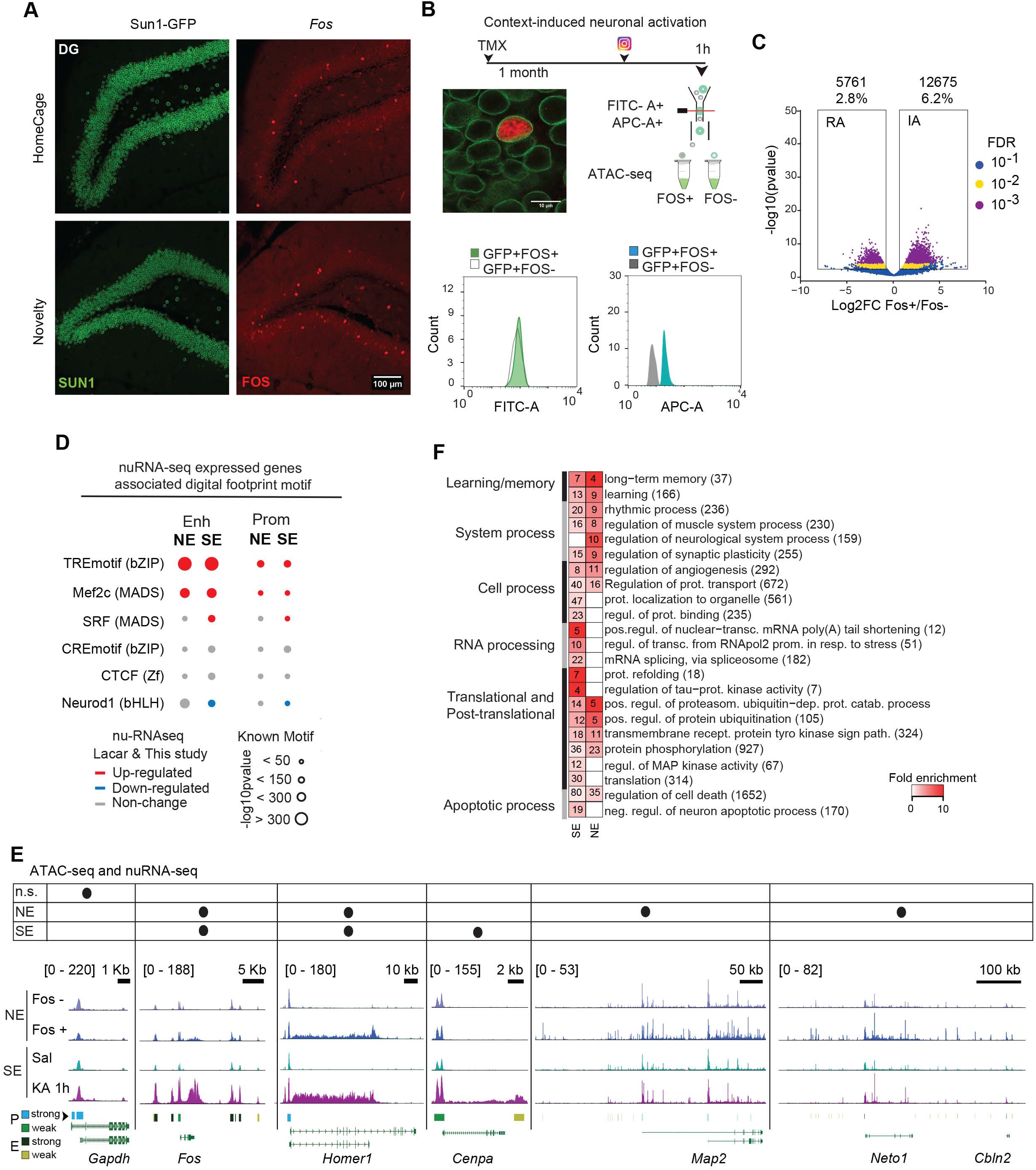
Physiological and pathological neuronal activation share common mechanisms but trigger distinct genomic signatures. **A**. Confocal images of granular neurons in the dentate gyrus (DG) of mice maintained in their homecages or exposed to 1 h of novelty exploration (NE). Brain slices were stained against GFP (green) and Fos (red). **B**. Experimental design of the approach used to study chromatin changes in neurons activated by 1 h of NE. Bottom: channel filtering of flow cytometry signal for Sun1-GFP+ (FITC-A) singlet nuclei and differences in Fos staining (APC-A). **C**. Volcano plot showing the significance value distribution after differential accessible regions analysis; upper values indicate number of regions and percentage of all the accessible regions detected. **D**. Digital footprinting at enhancer and promoter sites in NE and SE, indicating the motif enrichment (circle size) and the associated TF expression change in NE ^40^ and SE (this study) nuRNA-seq datasets (upregulated TFs are shown in red, downregulated one in blue, and those not changed in gray). **E**. Comparison of ATAC-seq tracks for physiologically (Fos^+^ *vs* Fos^−^ neurons) and chemically activated (Sal *vs* KA-1h) neurons, at the DEG in the SE (this study) and NE ^40^ nuRNAseq datasets. The table indicates the increase in accessibility at the gene body and nuRNA upregulation per each condition. Bottom bars indicate the regions classified as strong or weak at promoters and enhancers. Values indicate the levels of counts in RPM. **F**. Heatmap of fold enrichment for biological process GO terms in genes displaying IA in the NE and SE datasets. Values correspond to the number of genes associated with each term. See also Figure S5.

Although most of the genes responding to NE at the chromatin accessibility and transcriptional levels were retrieved during SE, a subset of genes was unique to this paradigm (Fig. 5E). Consistently, GO enrichment analyses reveled a greater enrichment for neuronal plasticity and memory functions in the NE-specific set, while the SE-specific genes were related to the modification and processing of RNA and protein, including the regulation of Tau and kinases involved in neurophysiopathology (Fig. 5F). The comparison of chromatin accessibility profiles also revealed noticeable differences between SE and NE (**Fig. S5B**; comparison of **Figs. S5F** and **S4G**). For instance, the TF footprints detected at DARs exclusive to the NE situation do not show *de novo* RNAPII or CBP binding after SE (**Fig. S5G**). However, the same sites displayed reduced DNA methylation and enhanced H3K4me1 and H3K27ac (two HPTMs associated with enhancer activation) after fear conditioning in a new context ^41^ (**Fig. S5H**). These results underscore both the common mechanisms and the stimuli-dependent differences in the genomic signature of pathological versus physiological activation of principal neurons.

### Activity-driven transcription is associated with the formation of gene loops and stronger promoter-enhancer interactions

Our multi-omic analysis indicates that activity-driven transcription is associated with an increase in chromatin accessibility and RNAPII occupancy that extends over the gene limit and points to the participation of structural proteins that topologically delimit locus responsiveness. To test this hypothesis, we investigated the 3D chromatin architecture of activated neurons using Hi-C (**Fig. S6A**). This genome-wide and fully unbiased chromatin conformation capture technique enables the analysis of compartments, CTCF loops and gene loops ^42^. We combined our FANS procedure with Hi-C to generate the first draft of DNA interactions in the hippocampus of adult behaving mice both in the basal situation and during SE (Fig. 6A). We detect increased interactions with nearby ATAC-seq peaks at example loci with increased expression after KA treatment (Fig. 6B). To test this on a genome-wide scale, we used Fit-Hi-C to retrieve a list of significantly enriched pairwise intra-chromosomal interactions (Table S5). Our analyses revealed enhanced interactions between proximal extragenic DARs and the TSS of AI genes (Fig. 6C; see **Fig. S6B** for an example involving the IEG *Fos*). We then called CTCF loops using HiCCUPS ^43^, but did not detect significant activity-driven changes, suggesting that CTCF loops are largely invariant during neuronal activation (**Fig. S6C-D**). This lack of change is consistent with our TFBS enrichment and digital footprint analyses (Figs. 4C and **S4I**). We then evaluated the appearance of gene loops between the TSS and TTS of genes with high levels of nuRNA-seq signal, and found increased formation upon SE (Fig. 6C-D; see **Fig. S6E** for an example at *Bdnf*). These gene loops between the TSS and the TTS of strongly induced genes may be essential for supporting the fast transcriptional rate of these loci. Their detection, together with the robust increase in gene body accessibility and intragenic RNAPII signal (Fig. 3E), point to a continuous re-loading of the RNAPII complex at IEGs during SE.

**Figure 6.**
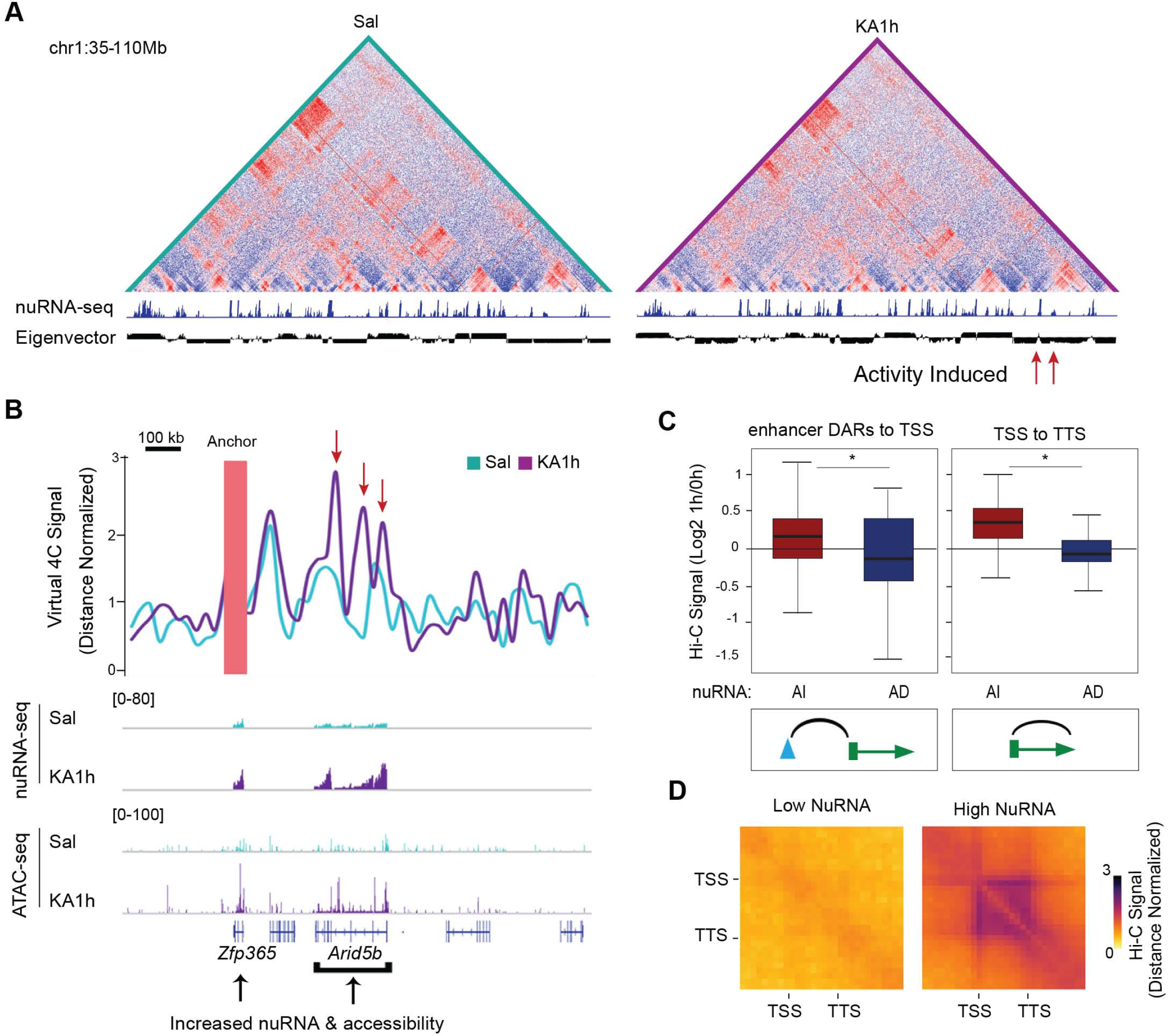
Neuronal activation induces gene loops and strengthens TSS-enhancer interactions. **A**. Hi-C maps of normalized distances in chromatin samples of saline and and KA-treated mice 1h after drug administration. Note the segregation of active and inactive chromatin in both situations, represented by positive and negative eigenvectors at the top and the augmented inflection (red arrow) in an SE-induced locus. **B**. Snapshot of a locus presenting strong *de novo* 4C-interactions in response to SE. **C**. Left: Changes to gene loop signal (TSS-TTS) 1 h after KA for activity-induced (AI, red) an activity-depleted (AD, blue) genes in nuRNA-seq. Right: Changes to Fit-Hi-C interaction signal 1 h after KA between differential ATAC-seq peaks and TSSs of genes with AI (red) or AD (blue) nuRNA-seq signal. AI and AD genes show increased and decreased gene looping interactions, respectively (*p<0.05 Wilcoxon rank sum test). **D**. Metaplot of Hi-C interactions at genes with low or high nuRNA-seq signal (Hi-C signal normalized by distance). See also Figure S6.

### Longitudinal analyses unveil chromatin changes that remain long after SE

Next, we investigated the duration and reversibility of transcriptional and chromatin accessibility changes by expanding our study to 6 h and 48 h after SE. In the case of nuRNA-seq (**Table S6**), the longitudinal analysis revealed a dynamic scenario in which the broad initial transcriptional response led to a secondary wave of changes that still affected thousands of transcripts but with more moderate changes, and with the transcriptome being largely restored to the basal situation 2 days later (Figs. 7A and **S7A-D**). Of note, there were more genes downregulated than upregulated at later time points, which contrasts with the oppositely skewed distribution observed during SE. Some of these downregulated genes are related to potassium and calcium transport, likely reflecting the late homeostatic response (**Fig. S7E**).

**Figure 7.**
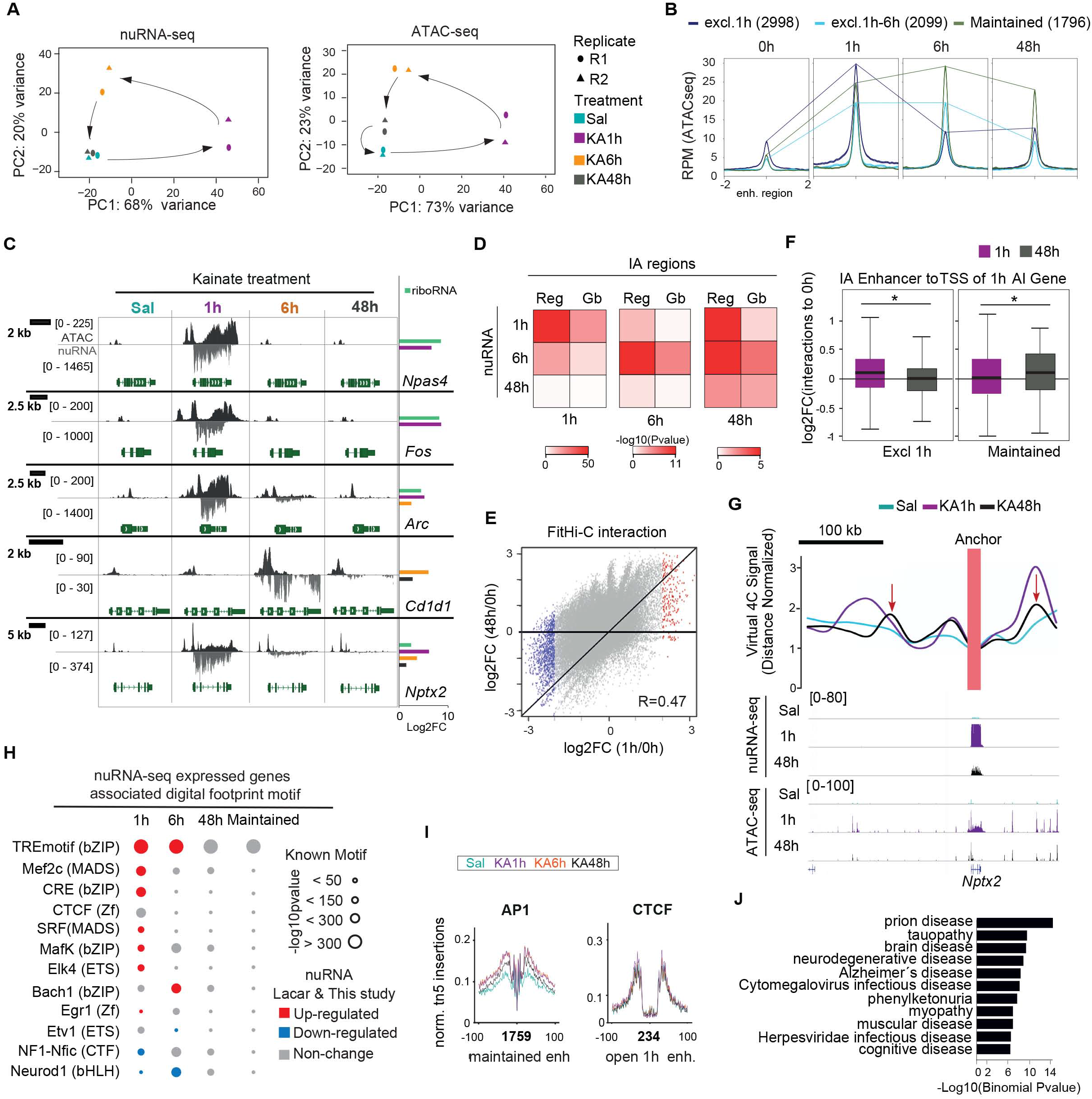
Long-lasting chromatin accessibility changes are associated with AP1 binding and disease. **A**. Principal component analysis (PCA) for nuRNA-seq (left) and ATAC-seq (right) datasets. Note that the Sal (0h) and KA-48h nuRNA-seq samples overlap distant from the KA-1h samples, while KA-6h samples are separated from both clusters confirming a second and independent wave of transcriptional changes. **B**. Comparison of the ATAC-seq signal at activity-regulated enhancers that show increased accessibility (IA) exclusively at 1 h, at 1 h and 6 h, or maintained for 48 h. **C**. Snapshots for ATAC-seq and nuRNA-seq tracks at the different time points. The examples illustrate the diversity of profiles under the general denomination of IEG. *Fos* and *Npas4* present a very rapid and transient induction associated with dramatic changes in the accessibility of the chromatin at the gene body, while other IEGs, such as *Arc* and *Nptx2*, maintain their induction for hours or even days after SE. Late response genes, such as *Cd1d1*, increase transcription and chromatin accessibility long after SE. Right bars show significant (FDR < 0.1) log2FC for nuclear transcripts at each time point. Values are shown in RPM. **D**. Heatmap showing the significance in the BETA analysis for changes in ATAC-seq and transcript levels in each time point at promoter/enhancer (Reg) or gene bodies (Gb). **E**. Total Fit-Hi-C interactions (gray) and their changes after 1 h (x-axis) or 48 h (y-axis) compared to 0 h. Dots correspond to the interactions that show more than a 4-fold increase (red) or decrease (blue) at 1 h. Most blue points are slightly below the horizontal, while most red points are above the horizontal, indicating that some of the SE-originated interactions are still maintained after 48 h. **F**. Changes to the Fit-Hi-C 1 h and 48 h interaction signals between IA exclusive 1h (left) and maintained (right) regions, and TSSs of activity-induced (AI) genes. AI genes show decreased and increased regulatory DARs interactions at 1 h and 48 h respectively (*p < 0.05 Wilcoxon rank sum test). **G**. Snapshots of the *Nptx2* and *Fos* loci presenting augmented interactions with an upstream enhancer in response to SE. This interaction remains slightly over the basal level even after 48 h. **H**. Digital footprinting at enhancer/promoter sites in SE dynamic, indicating the motif enrichment (circle size) and the associated TF expression change in NE nuRNA-seq datasets. Red: upregulated; blue: downregulated; gray: no-change. **I**. The plots show the footprinted sites at the AP1 and CTCF motifs detected at maintained and open at 1 h respectively, as well as the number of motifs detected (profiles indicate the normalized tn5 insertions). The colors indicate the comparison between 0 h, 1 h, 6 h, and 48 h after KA datasets. **J**. GREAT analysis reveals the set of disease-related processes enriched at maintained regions in AI genes. sd: system disease, sn: system neoplasm. See also Figures S7 and S8.

In contrast, the longitudinal analysis of chromatin accessibility changes (**Table S7**) did not reveal the same restoration to the original state. The samples corresponding to 48 h after KA approached the control samples (Sal), but did not overlap (Fig. 7A). Indeed, a relatively large subset of DARs (> 10,000), particularly those located at intergenic regions, persists 48 h after SE (Figs. 7B-C and **S7F-I**). Moreover, although BETA revealed a clear correlation between augmented transcript production and accessibility for each time point (Fig. 7D), in the case of extragenic regulatory regions, IA regions at 48 h showed a better correlation with transcriptional changes at 1 h and 6 h than at 48 h. Hi-C maps are consistent with ATAC-seq profiles and also present a large but incomplete restoration of interactions 48 h after SE (Fig. 7E-F). Together these intriguing results support the notion of a genomic memory in the form of protein binding or architectural modifications that originate from past transcriptional activity. In some genes such as *Nptx2*, encoding a synaptic protein involved in excitatory synapse formation ^44^, the chromatin change was associated with enhanced transcription (**Fig. S8A**, left), and a slight strengthening of the interaction between the promoter and a proximal enhancer (Fig. 7G). In other genes, such as *Kfl4* encoding a TF involved in the response to nerve and brain injury ^45^, the late activity-dependent occupancy was not linked to any apparent change in transcription (**Fig. S8A**, right).

Digital footprint analysis at regions displaying long-lasting changes revealed a specific enrichment for AP1 binding, while the enrichment for other activity-regulated TFs such as SRF and CREB was only detected at earlier time points (Fig. 7H-I). Interestingly, the AP1 sites presenting long-lasting changes after SE also responded to novelty exploration (**Fig S8B**), further underscoring the biological relevance of these sites. This binding signal might correspond to Fos-Jun dimers or to other TFs of the same family that recognize the same motif. Although the activity-induced transcription of AP1 genes is highly transient (Fig 7H), at the protein level, Fos immunoreactivity persists even 6 h after SE (**Fig. S8C**). Therefore, we cannot discard that a reduced number of AP1 molecules induced during seizure could still remain bound at specific loci at later time points. Notably, the genomic regions displaying long-lasting changes are associated with genes involved in brain diseases related to protein accumulation (Fig 7J), thus linking these changes with pathological traits. Consistent with this notion, mice that underwent SE displayed both memory deficits (**Fig. S8D**) and impaired induction of *Fos* (**Fig. S8E**) when subjected to a novel object location memory task (which is known to be hippocampal dependent). The *Fos* locus is surrounded by several regions that display long-lasting changes in occupancy, including sustained AP1 binding at putative enhancers located close to the neighbor gene *Jdp2* (**Fig. S8F**), which encodes for a repressor of AP1. These results suggest that the “silent” changes retrieved in our longitudinal study might affect the future responsiveness of the neuron and thereby contribute to hippocampal dysfunction in the epileptic brain.

## Discussion

Here, we investigated the transcriptional and chromatin changes occurring in neuronal nuclei of adult behaving mice upon activation. To gain cell-type specificity, we genetically tagged the nuclei and polysomes of excitatory hippocampal neurons for their isolation and use in several sequencing methods. Subsequent multi-omics analyses revealed an unexpectedly broad and dynamic scenario with multiple levels of activity-dependent regulation. For instance, the profiling of nuclear transcripts widens the scope of conventional mRNA screens and improves temporal resolution. Thanks to these features we found that the robust induction of IEGs is followed by a transient shutdown of metabolism genes and delayed downregulation of genes involved in ion transport and synaptic transmission. The later changes may contribute to the homeostatic plasticity mechanisms that compensate for prolonged activation and stabilizes neuronal firing ^46^. Our nuRNA-seq screen also reveals the production of ecRNA in activity-induced genes that may finetune the activity of these loci and contribute to transcriptional memory ^19^. Furthermore, its comparison with translatome data identifies genes in which synaptic activity specifically regulates ribosome engagement.

Further integration of nuRNA-seq with ATAC-seq, RNAPII ChIP-seq and Hi-C maps obtained in the same activation paradigm demonstrates for the first time that the robust transcription of IEGs during SE relies on the formation of gene loops that bring together the TSS and the TTS and favor the continuous re-loading of the RNAPII complex. Such organization has been described for highly transcribed genes in yeast ^47, 48^ and *Drosophila* cells ^42^, but not in neurons. Our analyses also identified hundreds of *de novo* or strengthened promoter-enhancer interactions and thousands of TF binding events in the chromatin of hippocampal excitatory neurons upon activation. Although some of these changes are transcription-independent and rely on the post-translational modification of TFs (e.g., phosphorylation), many others, such as the AP1 binding detected after both physiological and pathological stimulation, are transcription-dependent. Interestingly, the comparison of chromatin changes associated with physiological and pathological neuronal activation demonstrated that, in addition to common mechanisms there are remarkable differences in the scope and magnitude of the changes. These large-scale and dynamic adjustments of genome topology likely contribute to the rapid and coordinated transcriptional response associated with neuronal activation in both paradigms.

Intriguingly, our longitudinal analyses also retrieved changes in chromatin occupancy that persist even 2 days after stimulation. In particular, AP1 occupancy is detected long after the transcriptional shutdown of the encoding loci. This result is in agreement with a previous observation of stable accessibility changes in granular neurons 24 h after electrical stimulation ^30^. Functional genomics analyses indicate that these long-lasting changes may relate to brain diseases. Consistent with this link, a recent study showed that disease-associated SNPs are often found within cis-regulatory elements and, in particular, those within AP1 motifs show reduced chromatin accessibility ^49^. In a broader context, the enduring changes in accessibility that are associated with *de novo* AP1 binding to distal regulatory regions of activity-induced genes and with enhanced promoter-enhancer interactions may influence future activity of the loci. AP1 has been recently shown to recruit the BAF chromatin-remodeling complex to enhancers to locally augment chromatin accessibility ^50^. Therefore, the robust induction of AP1 subunits in response to activity might produce an excess of proteins to guarantee the occupancy of these sites and the priming effect. Since many of these loci play important roles in regulating synaptic plasticity and excitability, the experience fingerprints that persist in the chromatin may act as a form of metaplasticity by influencing the future response of the neuron to the same or other stimuli.

## Acknowledgments

We thank Eloisa Herrera, Jose P. Lopez-Atalaya and Yijun Ruan for critical reading of the manuscript, and Román Olivares, Nuria Cascales, Antonio Caler and the personnel of the sequencing facility at the CRG (Barcelona, Spain) and HudsonAlpha (Alabama, USA) for technical assistance. JF-A and MTL-C are recipients of fellowships from the Spanish Ministry of Science and Innovation (MICINN). AB research is supported by grants SAF2017-87928-R and SEV-2017-0723 from MICINN co-financed by ERDF, PROMETEO/2016/026 from the Generalitat Valenciana, and RGP0039/2017 from the Human Frontiers Science Program Organization (HFSPO). MJR is supported by the NIH Pathway to Independence Award K99/R00 GM127671. VGC research is supported by the U.S. Public Health Service Award (R01) GM035463 from the NIH. The content is solely the responsibility of the authors and does not necessarily represent the official views of the National Institutes of Health. The Instituto de Neurociencias is a “Centre of Excellence Severo Ochoa”.

## Author contributions (using CREdiT taxonomy)

Conceptualization, J.F-A. and A.B.; Methodology, J.F-A., and A.B.; Software, J.F-A., and M.J.R.; Formal analysis, J.F-A., M.J.R., and M.T.L-C.; Investigation, J.F-A., M.L., and A.M.M-G.; Data Curation and Visualization, J.F-A., M.T.L-C.; Resources, B.dB.; Writing – Original Draft, A.B. and J.F-A.; Supervision, A.B. and V.G.C.; Funding Acquisition, A.B. and V.G.C.

