## Supplementary Methods and Figures for "Immediate and deferred epigenomic signature of neuronal activation"

### Inventory of Supplementary materials


- Supplemental Experimental Procedures and bibliography (pages 2-14)
- Supplemental figures S1-S8 and supplemental legends (pages 15-30)
- Supplementary Tables S1 to S8
    - **Table S1. Related to Figure 1.** DETs upon SE in nuRNA-seq analysis (see attached Excel file).
    - **Table S2. Related to Figure 2.** DTGs upon SE in riboRNA-seq analysis (see attached Excel file).
    - **Tables S3A-C. Related to Figure 3.** DARs upon SE and TPL treatment retrieved in the ATAC-seq analysis (see attached Excel file).
    - **Table S4. Related to Figure 5.** DARs upon NE in ATAC-seq physiological analysis (see attached Excel file).
    - **Table S5. Related to Figure 6.** Fit-Hi-C table (see attached Excel file).
    - **Table S6. Related to Figure 7.** DETs upon SE in the longitudinal nuRNA-seq analysis (see attached Excel file).
    - **Table S7. Related to Figure 7.** DARs upon SE in the longitudinal ATAC-seq analysis (see attached Excel file).
    - **Table S8. Related to Methods.** Information related to NGS data (see attached Excel file).

**Supplemental Experimental Procedures**

*Animals and treatments*

The generation of CaMKIIα-creERT2 (Erdmann et al., 2007) (from the European Mouse Mutant Archive EMMA *s*train 02125), the cre-dependent riboTRAP (Stanley et al., 2013) (Jackson labs, stock #030305) and Sun-1 tagged mice (Mo et al., 2015) (Jackson labs, stock #021039) have been previously described. Mice were maintained and bred under standard conditions, consistent with Spanish and European regulations and approved by the Institutional Animal Care and Use Committee. All strains where maintained in a pure C57BL/6J background. Experiments were conducted in 3-6 months old male animals. Mice received intragastric administration of tamoxifen as described in (Fiorenza et al., 2016) when they were ~2 month-old, and we wait 3-4 weeks before experimentation. For the induction of SE by KA (Milestone PharmTech USA Inc.), a concentration of 25 mg/kg was injected intraperitoneally. For the blocking of transcription by TPL (Triptolide from Tripterygium wilfordii, Abcam ab120720) a concentration of 0.8 mg/kg was injected intraperitoneally per mice 2 h before the injection of KA or Saline.

*Behavioral tasks*

For novelty exposure (NE), individually housed mice received 3-4 days of 5 min handling before exposing them to a novel environment. The arena consisted in a white acrylic box of 48 x 48 x 30 cm containing objects with different forms and colors. A total of six mice were individually placed during 60 min before microdissection of the hippocampus and FANS. The novel object location memory test followed the protocol of (Vogel-Ciernia et al., 2013). Briefly, one week after saline or KA administration, a total of 8-9 mice per condition were individually habituated to an empty arena (2 exposures of 5 min in consecutive days). For training, the same mice were placed for 10 min in the arena with two identical objects 24h after the last habituation session. For location memory testing 24 h after training, one of the two objects was moved to a new location. The time spent exploring each object was quantified and a Differential Index (DI = time exploring the displaced object – time exploring the non-displaced object / total time exploring both objects) was calculated.

*Immunohistochemistry and microscopy*

Mice were anesthetized with intraperitoneal injection of ketamine/xylazine, perfused with 4% paraformaldehyde in PBS and extracted brains were postfixed O/N at 4ºC. Coronal or sagittal vibratome 40 µm sections were obtained, washed in PBS and PBS−0.1% Triton X-100 (PBT). Sections were incubated 30 min with blocking buffer 5% newborn calf serum (NCS, Sigma N4762) – PBT, and incubated O/N at 4ºC with the primary antibodies diluted in 5% NCS-PBT. The primary antibodies used in this study are α -NeuN (1:2000, Chemicon MAB377), α -GAD67 (1:800, Millipore MAB5406), α-Fos (1:500, Synaptic Systems 226004) and α-GFP (1:1000, Aves Labs GFP-1020). After primary antibody incubation and washing with PBS, sections were incubated with Alexa Flour conjugated IgG (1:500, Life Technologies) for fluorescent labeling. Nuclei were counterstained with a 1 nM DAPI solution (Invitrogen) before mounting with Fluoromount (Sigma). Images were taken using Olympus confocal inverted microscope and processed using ImageJ.

*Ribosomal-RNA immunoprecipitation sequencing*

TRAP immunoprecipitation followed the original protocol by (Heiman et al., 2014) with small changes in the extraction step. Hippocampi were micro-dissected after cervical dislocation, washed in PBS and submerged into ribosomal extraction buffer (REB-LOW: 150 mM KCl, 10 mM $MgCl_2$, 20 mM Hepes-KOH (pH 7.8), 1% IGEPAL

CA-630, 0.5 mM β-mercaptoethanol, 1x proteinase inhibitors (cOmplete EDTA-free, Roche), 60 U/ml RNasin Plus RNasa (Promega N2611), 100 μg/ml cycloheximida). Tissue was homogenized (25 times with pestle A and B) using a dounce (K8853000002, Kontes-DWK). A total of six hippocampi were pooled together and nuclear supernatant was obtained by centrifugation at 2000xg 4ºC for 10 min. Next steps followed the original protocol.

*Fluorescent Activated Nuclei Sorting (FANS)*

Novel protocol was designed for the maximum efficiency and specificity, a total of 1M nuclei/mice. Hippocampi were microdisected after cervical dislocation, rapidly washed in PBS and extracted with the dounce homogenizer (Sigma) with 1ml of Nuclear extraction buffer (NEB: 250 mM sucrose, 25 mM KCl, 5 mM $MgCl_2$, 20 mM Hepes-KOH (pH 7.8), 0.5% IGEPAL CA-630, 65 mM β-gycerol, 1mM β-mercaptoethanol, 1x proteinase inhibitors (Roche, cOmplete EDTA-free), 0.2 mM spermine, 0.5 mM spermidine, 60 U/ml RNasin Plus RNasa (Promega N2611), 5% NCS). Tissue was disrupted 15 times with pestle A and B, as described above. In case of nuclei fixation, formaldehyde was added in a final concentration of 1% after extraction, incubated at room temperature in rotation for 10', and blocked in rotation with 1,25 mM gycline. Hippocampi from 3 (SE experiments) to 6 (NE experiment) mice were pooled during filtration in a 35 μm mesh capped tube. In case of nuclear staining the nuclei pool was split in two tubes of 750 μl total NEB and were stained during 30 min with the primary antibody α-Fos-Alexa647 (1:40, sc-52 AF647) or α-NeuN (1:100, Chemicon MAB377) at 4ºC in rotation. In case of α-NeuN, the secondary antibody Alexa Fluor-conjugated IgG (1:1000, Life Technologies) was incubated during 15' more at 4ºC in rotation. Nuclei were stained 10 min with DAPI at a final concentration of 0.01 mM. After incubation, the nuclei preparation was diluted in Optiprep (O) density gradient medium (60% iodixianol, Sigma D1556) up to a concentration of 22%. A density gradient was prepared in 13 ml ultracentrifugue tube containing at the bottom a layer of O-44% and an upper layer of O-22% with the nuclear extraction. The tube was centrifuged at 7,500 rpm for 23 min at 4ºC and nuclear fraction was collected and transferred into a tube with the same volume of nuclear isolation buffer (NIB: 340 mM sucrose, 25 mM KCl, 5 mM $MgCl_2$, 20mM Hepes-KOH (pH 7.8), 65 mM β-glycerol, 1x proteinase inhibitors (cOmplete EDTA-free, Roche), 0.2 mM spermine, 0.5 mM spermidine, 60 U/ml RNasin Plus RNasa (Promega N2611), 5% NCS). Nuclei populations were selected with FACS Aria2, selecting DAPI staining, singlet nuclei, Sun1-GFP positive, and sorted into a tube with NIB. Flow cytometry data was analyzed and plot in *Flowjo*.

*RT-qPCR and RNA library construction*

Bulk RNA extraction from dissected hippocampi was done with TRI reagent (Sigma-Aldrich). For TRAPped RNA, after O/N incubation, beads were washed 3-4 times in ribosomal extraction buffer (RIB-HIGH: 350 mM KCl, 10 mM $MgCl_2$, 20 mM Hepes-KOH (pH 7.8), 1% IGEPAL CA-630, 0.5 mM β-mercaptoethanol) and re-suspended in 350 μl of RLT for purification with RNeasy Micro kit (Qiagen 74004). For nuclear RNA, after FANS, 2 millions nuclei were pelleted at 4ºC 1,000 g for 7 min and re-suspended in RLT for purification with RNeasy Micro kit (Qiagen 74004). qPCR was performed in Applied Biosystems 7300 real-time PCR unit. RNA samples were reverse transcribed using RevertAid First-Strand cDNA syntesis kit (Fermentas) and quantified using Eva Green qPCR reagent mix. Samples were assayed in duplicates and the results were normalized to *Gapdh* transcript levels. The sequences of the primers used in RT-qPCR assays are:

| | |
|---|---|
| Fos Fw | 5´-GCTTCCCAGAGGAGATGTCTGT |

| | |
|---|---|
| Fos Rv | 5´-GCAGACCTCCAGTCAAATCCA |
| Npas4 Fw | 5´-CTGGCCCAAGCTTCTTCTCA |
| Npas4 Rv | 5´-TCCATGCTTGGCTTGAAGTCT |
| Arc Fw | 5´-GCAGGAGAACTGCCTGAACAG |
| Arc Rv | 5´-AAGACTGATATTGCTGAGCCTCAA |
| Plp1 Fw | 5'-CTTTGGCGACTACAAGACCAC |
| Plp1 Rv | 5'-ACAGTCAGGGCATAGGTGATG |
| Gfap Fw | 5´-GGACAACTTTGCACAGGACCTC |
| Gfap Rv | 5´-TCCAAATCCACACGAGCCA |
| Bdnf Fw | 5´-GAAGGTTCGGCCCAACGA |
| Bdnf Rv | 5´-CCAGCAGAAAGAGTAGAGGAGGC |
| Gapdh Fw | 5’-CATGGACTGTGGTCATGAGCC |
| Gapdh Rv | 5’-CTTCACCACCATGGAGAAGGC |

RNA-seq, library preparation and sequencing were performed at facilities in the Center for Genomic Regulation (CRG, Barcelona, Spain). Truseq libraries were prepared from ribo-depleted total RNA in the case of nuRNA-seq and from polyA-selected mRNA in the case of riboRNA-seq. Samples were sequenced (single-end, 50 bp length) using a Illumina HiSeq2500 apparatus with a depth of at least 80M reads in nuRNA and 30M reads in riboRNA (**Table S8**).

*ATAC assay and library construction*

After FANS 50K nuclei were pelleted at 4ºC 1,000 g for 7 min, re-suspended in 50 μl of transposition reaction (Illumina FC-121-1030) and gently pipette to resuspend the nuclei. The assay of transposase-accessibility chromatin coupled to massive sequencing (ATAC-seq) experiments were conducted as the original protocol (Buenrostro et al., 2013). Briefly, nuclei were incubated with tn5 during 30 min at 37ºC, and immediately purified with MinElute PCR purification kit (Qiagen 28004). In order to reduce PCR induced biases during library preparation, after the first 5 PCR cycles, 10% of the sample was monitored with SYBR-green qPCR for 20 cycles to calculate the 25% of saturation per sample (any sample needed more than 12 PCR cycles in total). Libraries were purified with MinElute PCR purification kit (Qiagen 28004). ATAC-assay was performed in an Applied Biosystems 7300 real-time PCR unit using Eva Green qPCR reagent mix. Samples were assayed in duplicates and results were normalized to Gapdh promoter insertion levels. The sequences of the primer pairs used in ATAC-qPCR assays are:

| | |
|---|---|
| Heterochromatin Fw | 5´-CTACCGAGTGTTGATTGCCGT |
| Heterochromatin Rv | 5´-TGATGCAAGTGTCAAGCTCAATG |
| Fos-promoter Fw | 5´-GCAGTCGCGGTTGGAGTAGT |
| Fos-promoter Rv | 5´-CGCCCAGTGACGTAGGAAGT |
| Gfap-promoter Fw | 5´-TACCAGAAAGGGGGTTCCTT |
| Gfap-promoter Rv | 5´-AACTCCTCTCACCCCACTGA |
| Gapdh-promoter Fw | 5’-TTCACCTGGCACTGCACAA |
| Gapdh-promoter Rv | 5’-CCACCATCCGGGTTCCTATAA |

For ATAC-seq, sequencing (paired-end, 50bp length) was performed at the CRG facilities (Barcelona, Spain) using a Illumina HiSeq2500 apparatus and a depth of at least 100M reads per sample (**Table S8**).

*ChIP- assay and ChIP library construction*

For Bulk ChIP the whole hippocampi from two mice were microdissected and pooled together. For sorted ChIP after FANS, 2 millions nuclei were pelleted at 4ºC 2,000 g for 7 min and re-suspended in SDS-lysis buffer or RIPA-PBS buffer. ChIP was conducted as previously described (Lopez-Atalaya et al., 2013) using the anti-H3K4me3 (Millipore 07-473), anti-CBP (sc583x) and anti-RNAPII (sc901x). ChIP-assay was performed in an Applied Biosystems 7300 real-time PCR unit using Eva Green qPCR reagent mix. Samples were assayed in duplicates and results were normalized to input. The sequences of the primer pairs used in ChIP-qPCR assays are:

| | |
|---|---|
| Camk2a-promoter Fw | 5´-GAGTCCTAGAGAGCATGGGG |
| Camk2a-promoter Rv | 5´-TGCATACACAGTGCTTCCAAA |

| | |
|---|---|
| Heterochromatin Fw | 5´-CTACCGAGTGTTGATTGCCGT |
| Heterochromatin Rv | 5´-TGATGCAAGTGTCAAGCTCAATG |

For ChIP-seq, library preparation and sequencing (single-end, 50bp length) were performed at the CRG facilities (Barcelona, Spain) in a Illumina Hi2500Seq apparatus with a depth of 40M reads per sample (**Table S8**).

*In situ Hi-C library construction*

After FANS 2M nuclei were pelleted at 4ºC 2,000 g for 10 min. The nuclei were resuspended in ice-cold HiC lysis buffer and the original in situ Hi-C protocol by (Rao et al., 2014) was followed. Libraries were prepared with TruSeq and sequenced (paired-end 50bp) with Illumina NovaSeq apparatus at the Hudson Alpha sequencing facility (Alabama, AL, USA) with a depth of 1B reads per sample (**Table S8**).

*Genomic data processing and access*

All sequenced datasets adapters were trimmed using *cutadapt* v1.18 (Martin, 2011) and aligned to mm10. Only reads with mapq > 30 and mapping to nuclear chromosomes were used for the posterior analysis. Data was processed with the extensive use of custom scripts and Samtools (v1.9) (Li et al., 2009), bedtools v2.26.0 (Quinlan and Hall, 2010) and DeepTools v3.1.0 (Ramirez et al., 2016). Whole genome alignments were normalized to RPM (read per million sequenced reads) and visualized using IGV (v2.3.92) (Thorvaldsdottir et al., 2013). Data can be accessed at the GEO repository using the accession number GSE125068.

*RNA-seq data*: Reads were aligned with HISAT2 v2.1.0 (Kim et al., 2015). For a clear comparison between riboRNA and nuRNA datasets, reads were annotated to exons from Ensemble (GRCm38.89) and quantified using Rsubreads (Liao et al., 2014). Differential expression analysis was performed in DESeq2 (v1.10.0) (Love et al.,

2014) counting for batch and treatment effect. Genes with FDR<0.1 and log2FC > +/-1 were considered significantly changed. Transcriptome analyses used the datasets GSE74971 (Halder et al., 2016) and GSE77067 (Lacar et al., 2016) applying the same cut-offs used in our study. GO terms where analyzed using pantherdb.org with Fisher's Exact with FDR multiple test correction, cut-off was done by the number of genes associated to category (< 2,000 genes), the number of DEG associated to the category (> 3 genes) and fold enrichment > 3 (p-value < 0.05). Over-Representation Enrichment Analysis (ORA) was performed with WebGestalt, and the use of the DisGeNET human database (Bauer-Mehren et al., 2010; Queralt-Rosinach et al., 2016).

*ATAC-seq data:* Reads were aligned with bowtie2 v2.2.6 (Langmead and Salzberg, 2012). Duplicated reads were removed with Picardtools and only paired reads were used for the posterior analysis (the KA-6h biological replicate 1 had to be down-sampled). The same pipeline was applied to ATAC-seq datasets from (GSE63137, Mo et al., 2015) and (GSE82015, Su et al., 2017). Peak calling was done with MACS2 (Zhang et al., 2008) in the merged samples that were going to be compared for Differential Accessible Regions (DARs), quantified using Rsubreads on called peaks with FDR < 1e-5. For Differential Accessible Genes (DAGs) reads were annotated to the whole gene region from Ensembl (GRCm38.89), quantified using Rsubreads (Liao et al., 2014). DARs and DAGs analysis was performed using DESeq2 (v1.10.0) (Love et al., 2014) counting for batch and treatment effect. Regions or Genes with FDR<0.1 and log2FC > +/-1 were considered significantly changed. The analysis of changes in TPL and NE experiments, were performed in the DESeq2 including in the experiment design the comparisons with KA-1h and Saline respectively that support the reduced n for the rlog calculation and value transformations. Unfortunately, due to the broad increase of accessibility at the gene bodies, we could not subtract polymerase driven transcriptional events from putative enhancers located at introns; the enhancer regions considered in this study were those positioned a more than 1 Kb from the TSS (upstream) or TTS (downstream) of annotated Ensembl genes (GRCm38.89). GO terms enrichment for DAGs were analyzed using pantherdb.org with Fisher's Exact with FDR multiple test correction, cut off was done by the number of genes associated to category (< 2000 genes), the number of DEG associated to the category (> 3 genes) and fold enrichment > 3 (p-value < 0.05). Disease ontology from genes associated to DARs was done with GREAT (McLean et al., 2010) using standard parameters. Promoter and enhancers regions were clustered with unsupervised method (k-means with linear normalization method) using seqMiner v1.3.4 (Ye et al., 2011). The *phastCons* score was obtained from the multiple alignments of 59 vertebrate genomes (60way Placental) to the mouse genome mm10 (Pollard et al., 2010; Siepel et al., 2005). Motif analysis was done at the promoter and enhancer sites with HOMER using validated database from ChIP-seq experiments. Motifs were classified by families (i.e. TRE motif is recognized by a number of proteins that form the AP1 complex) and by the detection of the related protein coding genes in our nuRNA-seq datasets. Predictive analysis of DARs and expression changes was done using BETA v1.0.7 basic activating/repressive function prediction with the Ensembl reference (GRCm38.89) and DEG from our nuRNA-seq datasets. Nucleosome occupancy information was retrieved with the *NucleoATAC* algorithm (Schep et al., 2015); the shifting analysis was conducted using custom R scripts 500 bp around the TSS. Digital footprint was done in DARs promoters/enhancers using the adapted *Wellington* algorithm for tn5

(Piper et al., 2013). ATAC-seq datasets from KA-1h were used to detect the footprints during SE (FDR < 0.01) and in the longitudinal analysis (p-value < 1e-10).

*ChIP-seq data:* Reads were aligned with bowtie2 v2.2.6 (Langmead and Salzberg, 2012). Chip-seq meta-analyses used data generated by (Halder et al., 2016; Kim et al., 2010; Malik et al., 2014; Scandaglia et al., 2017); in some cases the positions had to be trans-mapped to mm10. Peak calling was done with MACS2 v2.1.1 (Zhang et al., 2008) with default parameters.

*HiC data:* Hi-C datasets were processed using the Juicer pipeline (Durand et al., 2016) to align reads to the mm10 genome, filter duplicates, perform matrix normalizations, and obtain the eigenvector. Reads with a mapping quality >= 30 and binned at 25 kb were used in all downstream analyses. Distance normalization was done using the formula $\frac{(observed-expected)}{(expected+1)}$. Significant enhancer -promoter interactions were called using Fit-Hi-C (Ay et al., 2014) separately for each dataset with an FDR of 0.05. Fit-Hi-C interactions were compared across samples using the distance normalized signal. Intragenic interactions were visualized across genes with high and low nucRNA-seq signal categorized by the top and bottom quartiles respectively. High intensity interactions known as CTCF loops were identified via HiCCUPS (Rao et al., 2014) in each dataset and differential loops were determined by 2-fold changes in interaction signal in a combined total list of loops. Virtual 4C was derived by isolating all Hi-C interaction signal with each specified anchor. Because *Nptx2* lied at the border of two 25 kb bins, the average interaction signal across both bins was used.

*Statistical analyses*

All statistical analyses were two-tailed. For pairwise comparison of averages, data were tested for normality using Shapiro’s test. If any of the two samples were significantly non-normal, a non-parametric Mann-Whitney U/Wilcoxon rank-sum test was executed instead. When variances between groups were the same, a 2-way ANOVA was applied in the riboRNA-qPCR experiments. For multiple comparisons Kruskal-Wallis test was applied and Dunn test when the number of observations were unequal between groups. Bonferroni-corrected pairwise tests were used where appropriate post-hoc to correct for multiple comparisons. P values were considered to be significant when $\alpha < 0.05$. Mean ± s.e.m. Assuming the linear relation of the data, correlation was used. Bar plots and cumulative plots centered regions indicate the mean ± s.e.m, unless otherwise indicated. Heat maps show Euclidean clusters.

**Supplementary Figure legends**

**Supp. Figure S1. KA-induced neuronal activation causes broad changes in the nuclear transcriptome. A.** Step by step analysis and channel filtering (percentage of the filtered population indicated within the channel) of flow cytometry signal for the specific isolation of florescent singlet nuclei. Shown the comparison of intrinsic fluorescence in Sun1-GFP$^{-}$ (above) *vs* Sun1-GFP$^{+}$ (below) mice. **B.** Images of isolated nuclei immunostained with antibodies against GFP and the neuronal marker NeuN. **C.** Comparison of nuclear size of Sun1- GFP$^{+}$ (green) and wildtype nuclei stained with NeuN (pink). **D.** RT-qPCR in nuclear RNA showing Fos transcript levels in GFP positive nuclei after KA-induced SE (levels referred to GAPDH expression). **E.** ChIP-qPCR assay analysis for H3K4me3 in GFP positive and negative nuclei showing FANS specificity (values are normalized to input). Pr: promoter; HChrom: Heterochromatin. **F.** ATAC-qPCR assay showing specificity and promoter changes 1 h after KA-induced neuronal activation in GFP positive nuclei (levels referred to the *Gapdh* promoter). Pr: promoter; HChrom: Heterochromatin. **G.** Pearson correlation matrix between normalized samples clustered by Euclidian dendrogram. **H.** Percentage of nuRNA-seq reads aligned into exons. For comparison, we also present the values for riboRNA-seq samples. **I.** GO enrichment analysis for protein-coding DETs that are activity-induced (AI, red bars) or activity-depleted (AD, blue bars) by SE.

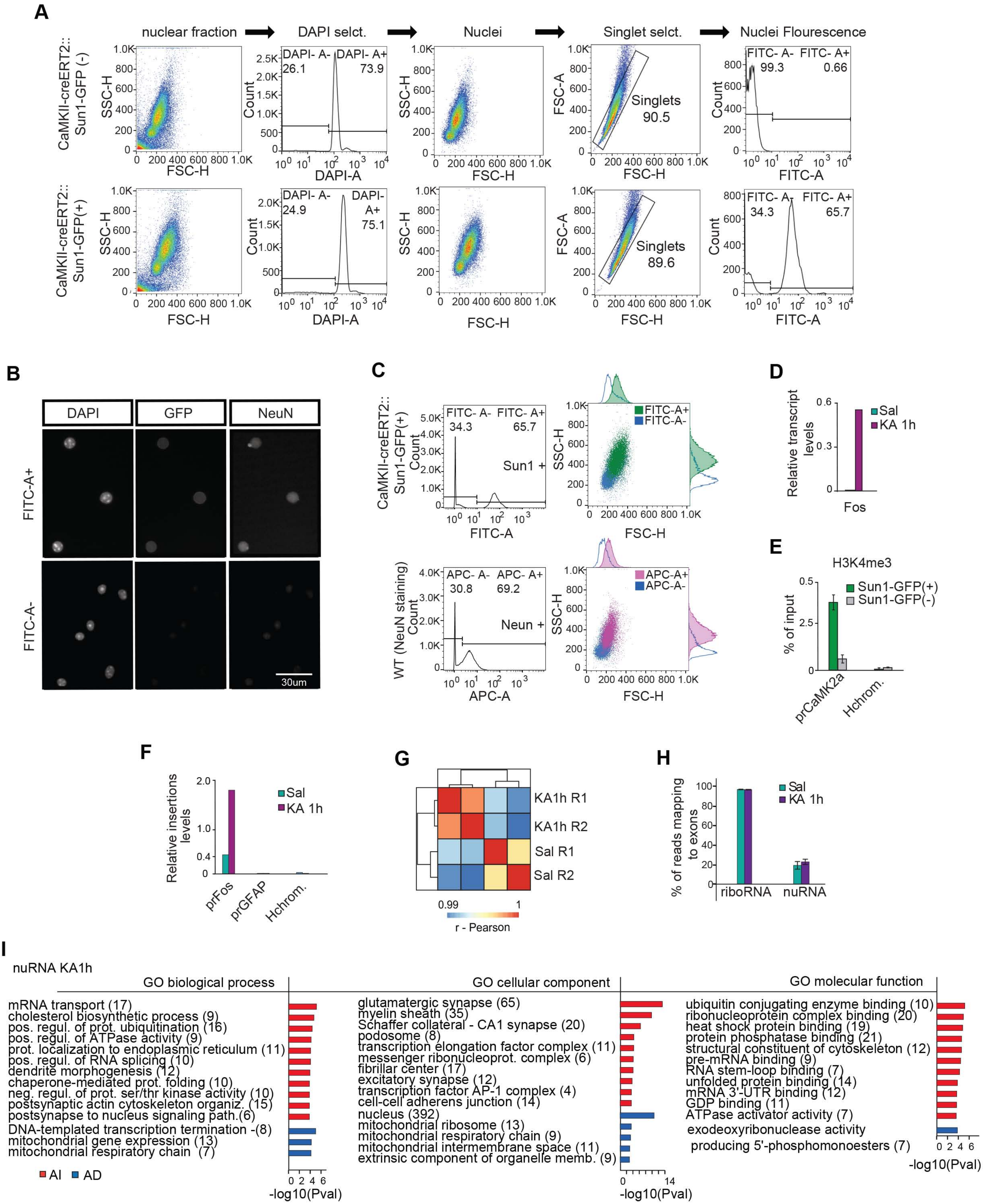
A
nuclear fraction
DAPI selct.
Nuclei
Singlet selct.
Nuclei Flourescence
CaMKII-creERT2::
Sun1-GFP (-)
SSC-H
FSC-H
DAPI- A-
26.1
DAPI- A+
73.9
Count
DAPI-A
FSC-A
Singlets
90.5
FITC- A-
99.3
FITC- A+
0.66
FITC-A
CaMKII-creERT2::
Sun1-GFP(+)
DAPI- A-
24.9
DAPI-
A+
75.1
Singlets
89.6
FITC- A-
34.3
FITC- A+
65.7
B
DAPI
GFP
NeuN
FITC-A+
FITC-A-
30um
C
CaMKII-creERT2::
Sun1-GFP(+)
Count
FITC- A-
34.3
FITC- A+
65.7
Sun1 +
FITC-A
FITC-A+
FITC-A-
SSC-H
FSC-H
WT (NeuN staining)
APC- A-
30.8
APC- A+
69.2
Neun +
APC-A
APC-A+
APC-A-
D
Relative transcript
levels
Sal
KA 1h
Fos
E
H3K4me3
% of input
Sun1-GFP(+)
Sun1-GFP(-)
prCaMK2a
Hchrom.
F
Relative insertions
levels
Sal
KA 1h
prFos
prGFAP
Hchrom.
G
KA1h R1
KA1h R2
Sal R1
Sal R2
0.99
1
r - Pearson
H
% of reads mapping
to exons
Sal
KA 1h
riboRNA
nuRNA
I
nuRNA KA1h
GO biological process
mRNA transport (17)
cholesterol biosynthetic process (9)
pos. regul. of prot. ubiquitination (16)
pos. regul. of ATPase activity (9)
prot. localization to endoplasmic reticulum (11)
pos. regul. of RNA splicing (10)
dendrite morphogenesis (12)
chaperone-mediated prot. folding (10)
neg. regul. of prot. ser/thr kinase activity (10)
postsynaptic actin cytoskeleton organiz. (15)
postsynapse to nucleus signaling path.(6)
DNA-templated transcription termination -(8)
mitochondrial gene expression (13)
mitochondrial respiratory chain (7)
AI
AD
-log10(Pval)
GO cellular component
glutamatergic synapse (65)
myelin sheath (35)
Schaffer collateral - CA1 synapse (20)
podosome (8)
transcription elongation factor complex (11)
messenger ribonucleoprot. complex (6)
fibrillar center (17)
excitatory synapse (12)
transcription factor AP-1 complex (4)
cell-cell adherens junction (14)
nucleus (392)
mitochondrial ribosome (13)
mitochondrial respiratory chain (9)
mitochondrial intermembrane space (11)
extrinsic component of organelle memb. (9)
-log10(Pval)
GO molecular function
ubiquitin conjugating enzyme binding (10)
ribonucleoprotein complex binding (20)
heat shock protein binding (19)
protein phosphatase binding (21)
structural constituent of cytoskeleton (12)
pre-mRNA binding (9)
RNA stem-loop binding (7)
unfolded protein binding (14)
mRNA 3'-UTR binding (12)
GDP binding (11)
ATPase activator activity (7)
exodeoxyribonuclease activity
producing 5'-phosphomonoesters (7)
-log10(Pval)

**Supp. Figure S2**. **SE-induced RNA translation activates nucleus-related functions**. **A.** Confocal images of principal hippocampal neurons in the DG and CA1 subfield of CaMKII-creERT2::GFP-L10a mice stained against GFP (green) and the interneuron marker GAD67 (red). Cells positive for GAD67 do not express GFP. **B.** RT-qPCR assay comparing transcript levels before IP (Bulk) and after immunoprecipitation (IP). Note that fold changes for IEG induction were larger after TRAPping. 2-way ANOVA p-values: * = treatment effect (KA 1h *vs* Sal); # = procedure effect (IP *vs* Bulk). */#: $p < 0.05$; **/##: $p < 0.01$; ***/###: $p < 0.001$; ns: non significant. **C.** Pearson correlation matrix between normalized riboRNA-seq samples clustered by Euclidian dendrogram. **D.** Principal component analysis of riboRNA-seq samples. **E.** Genomic snapshots for each replicate riboRNA-seq track at representative examples of non-neuronal (*Plp1*), housekeeping (*ActB*) and activity-induced (*Fos*) genes. Values indicate the number of counts in RPM. **F.** Gene biotype classification for DTGs 1h after KA. **G.** GO enrichment analysis for protein-coding genes displaying enhanced (red bars) or reduced (blue bars) translation upon SE. **H.** Comparison of nuRNAseq and riboRNAseq tracks for representative transcripts that are either detected as upregulated or downregulated in both screens (but showing largest changes in nuRNA-seq), or that show discordant changes in the two screens. Values indicate the levels of counts in RPM.

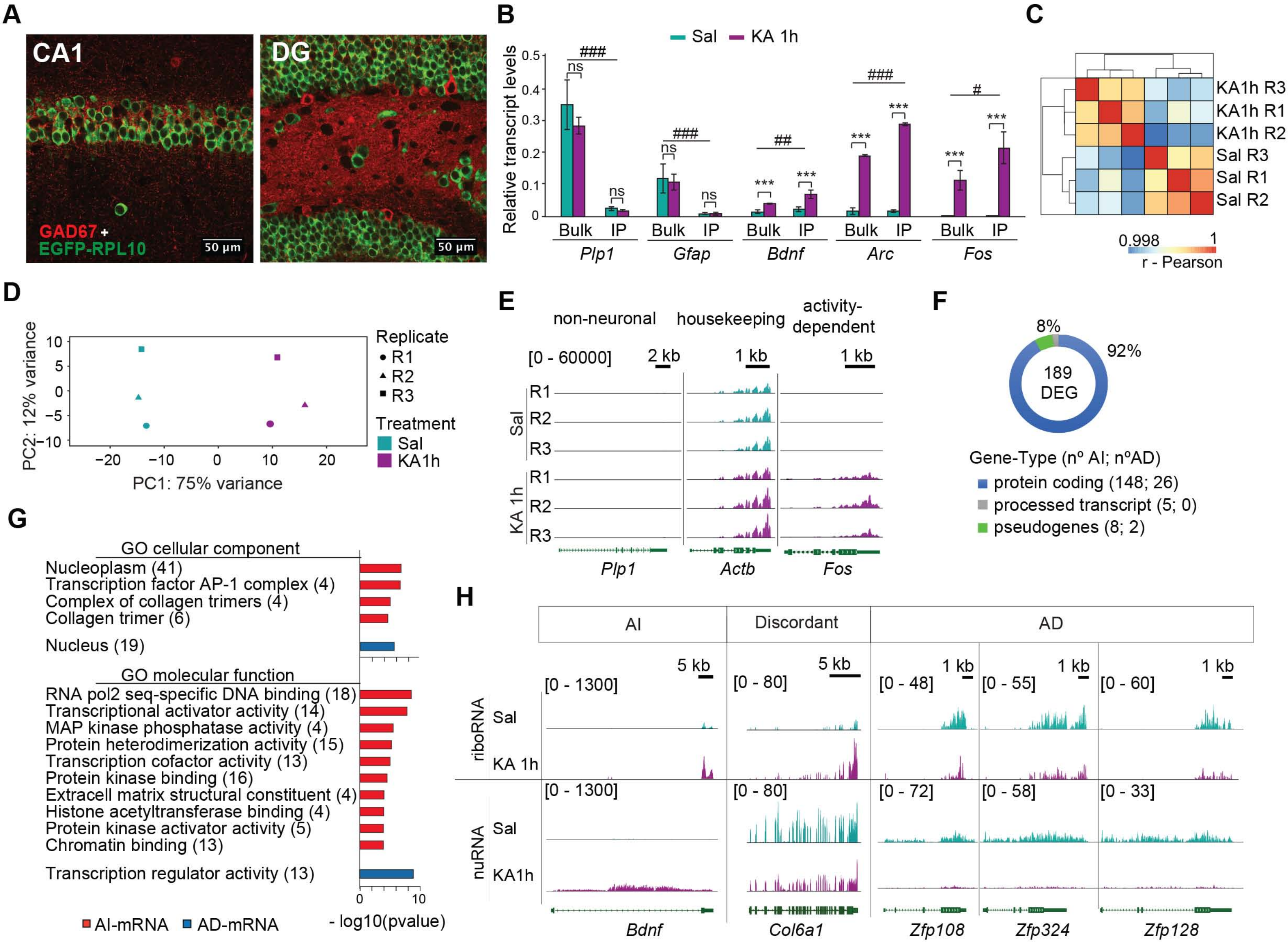
A
CA1
DG
GAD67 +
EGFP-RPL10
50 μm
B
Sal
KA 1h
Relative transcript levels
Bulk IP
Plp1
Gfap
Bdnf
Arc
Fos
C
KA1h R3
KA1h R1
KA1h R2
Sal R3
Sal R1
Sal R2
0.998
1
r - Pearson
D
PC1: 75% variance
PC2: 12% variance
Replicate
R1
R2
R3
Treatment
Sal
KA1h
E
non-neuronal
housekeeping
activity-dependent
[0 - 60000]
2 kb
1 kb
Actb
F
8%
92%
189 DEG
Gene-Type (nº AI; nºAD)
protein coding (148; 26)
processed transcript (5; 0)
pseudogenes (8; 2)
G
GO cellular component
Nucleoplasm (41)
Transcription factor AP-1 complex (4)
Complex of collagen trimers (4)
Collagen trimer (6)
Nucleus (19)
GO molecular function
RNA pol2 seq-specific DNA binding (18)
Transcriptional activator activity (14)
MAP kinase phosphatase activity (4)
Protein heterodimerization activity (15)
Transcription cofactor activity (13)
Protein kinase binding (16)
Extracell matrix structural constituent (4)
Histone acetyltransferase binding (4)
Protein kinase activator activity (5)
Chromatin binding (13)
Transcription regulator activity (13)
AI-mRNA
AD-mRNA
- log10(pvalue)
H
AI
Discordant
AD
riboRNA
nuRNA
5 kb
[0 - 1300]
[0 - 80]
[0 - 48]
[0 - 55]
[0 - 60]
[0 - 72]
[0 - 58]
[0 - 33]
Col6a1
Zfp108
Zfp324
Zfp128

**Supp. Figure S3. Transcriptional bursting increases accessibility at IEGs. A.** Fragment size distribution in ATAC-seq libraries. **B.** ATAC-seq signal at the TSS of highly expressed genes in NeuN$^+$ *vs* NeuN$^-$ CA1 cells (Halder et al., 2016). **C.** PCA of accessibility profiles shows similarity between cortical (Mo et al., 2015) and hippocampal excitatory neurons (this study). CTX_exc: cortical excitatory neurons; CTX_VIP: vasoactive intestinal peptide-expressing interneurons; CTX_PV: parvalbumin-expressing interneurons; HIPP_exc: hippocampal excitatory neurons. **D.** Percentage of reads mapped into nuclear and mitochondrial chromosomes in this study and (Su et al., 2017). **E.** Genome coverage in this study and (Su et al., 2017). E0 and E1: chromatin profiling coverage before (E0) and 1 h after synchronous electrical neuronal activation (E1). **F.** Comparison of genomic profiles at control and activity-regulated genes, including an extended view of the *Npas4* locus. Discrepancies may result from the superior genomic coverage and intrinsic differences between stimulation paradigms (chemical *vs.* electrical) and composition of the samples (hippocampal glutamatergic neurons nuclei *vs.* microdissected DG tissue). **G.** Left: Overlap between activity-induced genes detected in the nuRNA-seq and riboRNA-seq screens that show increased accessibility. Right: Heatmap representation of changes in accessibility and transcript levels in overlapping genes. **H.** Correlation between accessibility changes at gene bodies and riboRNA (left) and nucRNA (right) signal. Gray: genes retrieved in all the screens; blue: genes significantly regulated (FDR < 0.1). **I.** Metagene plot of RNAPII occupancy in AI genes (nuRNA-seq) split according to increased gene body accessibility (FDR > 0.1). **J.** Metagene plot of ATAC-seq signal in Sal, KA-1h and KA+TPL samples; genes were split as in **panel I**. **K.** TPL effect on the accessibility at gene bodies and TSSs of activity-regulated genes. **L.** Transcriptional bursting increases Tn5 insertions.

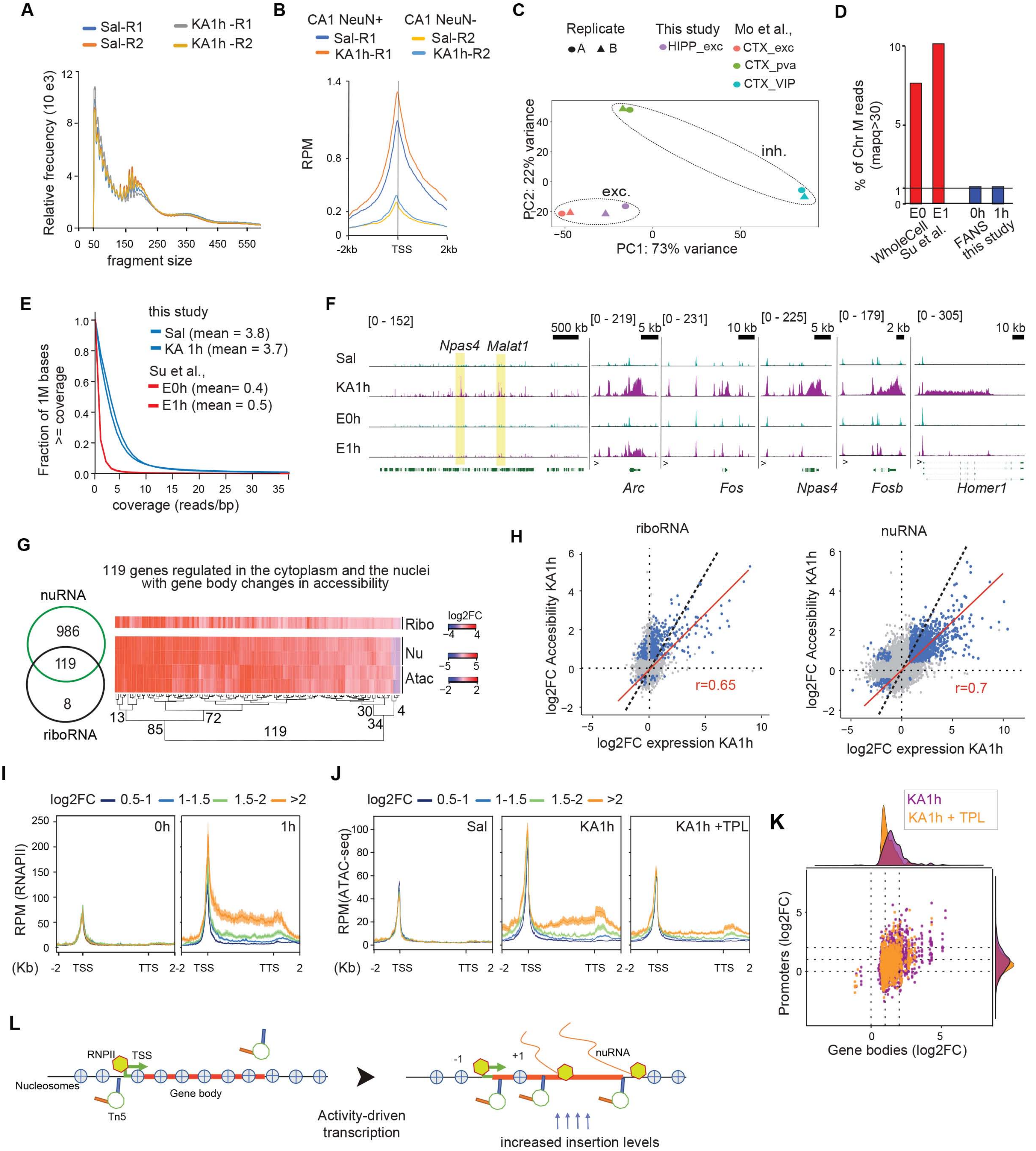
A
Sal-R1
Sal-R2
KA1h -R1
KA1h -R2
Relative frecuency (10 e3)
fragment size
B
CA1 NeuN+
CA1 NeuN-
Sal-R1
KA1h-R1
Sal-R2
KA1h-R2
RPM
-2kb
TSS
2kb
C
Replicate
A
B
This study
HIPP_exc
Mo et al.,
CTX_exc
CTX_pva
CTX_VIP
PC2: 22% variance
PC1: 73% variance
inh.
exc.
D
% of Chr M reads (mapq>30)
E0
E1
0h
1h
WholeCell Su et al.
FANS this study
E
this study
Sal (mean = 3.8)
KA 1h (mean = 3.7)
Su et al.,
E0h (mean= 0.4)
E1h (mean = 0.5)
Fraction of 1M bases >= coverage
coverage (reads/bp)
F
[0 - 152]
[0 - 219]
[0 - 231]
[0 - 225]
[0 - 179]
[0 - 305]
500 kb
5 kb
10 kb
5 kb
2 kb
10 kb
Npas4
Malat1
Sal
KA1h
E0h
E1h
Arc
Fos
Npas4
Fosb
Homer1
G
119 genes regulated in the cytoplasm and the nuclei with gene body changes in accessibility
nuRNA
986
119
8
riboRNA
Ribo
Nu
Atac
log2FC
13
72
85
119
30
4
34
H
riboRNA
nuRNA
log2FC Accesibility KA1h
log2FC expression KA1h
r=0.65
r=0.7
I
log2FC
0.5-1
1-1.5
1.5-2
>2
0h
1h
RPM (RNAPII)
(Kb)
TSS
TTS
J
log2FC
0.5-1
1-1.5
1.5-2
>2
Sal
KA1h
KA1h +TPL
RPM(ATAC-seq)
(Kb)
TSS
TTS
K
KA1h
KA1h + TPL
Promoters (log2FC)
Gene bodies (log2FC)
L
RNPII
TSS
Nucleosomes
Gene body
Tn5
Activity-driven transcription
-1
+1
nuRNA
increased insertion levels

**Supp. Figure S4. Activity-dependent TF-biding. A.** BETA analysis of accessibility changes at extragenic regions and associated changes in expression. IA: increased accessibility; RA: reduced accessibility. **B.** Bar plot comparing the percentages of DARs at accessible promoters and enhancers. Upper sector plots indicate percentage of strong/weak signal at IA and RA regions. **C**. Signal for RNAPII and CBP binding 1 h after KA. We compare IA and RA regions; IA regions are split between weak and strong promoters/enhancers. **D**. Signal of activity regulated profiles for H3K27ac 1 h after KCl (Malik et al., 2014) using the same classification than in **panel C**. Unt.: untreated. **E.** Signal for RNAPII and CBP 1h after KA in transcriptional-dependent and independent regions. **F.** Signal of H3K27ac 1 h after KCl (Malik et al., 2014) using the same classification than in **panel E**. **G**. Bar plot comparing the percentage of promoter/enhancer DARs annotated to nuRNA-seq regulated genes (FDR < 0.1). Upper sector plots show percentage of activity-induced (AI) and depleted (AD) genes associated with IA and RA regions. **H.** Genomic snapshot of ATAC-seq and RNAPII binding in saline, KA-1h and TPL+KA-1h samples at the *Npas4* locus (values in RPM). Annotations label the detected footprints (in red, less stringent footprints) and classification for the regions. Zoom-in inset shows upstream eRNA activity. **I.** Motif enrichment at transcription-dependent and -independent enhancers for the indicated activity-regulated TFs. **J.** Plots compare the digital footprint at AP1 and CTCF motifs in saline, KA-1h and TPL+KA-1h datasets (value correspond to normalized tn5 insertions). The bottom numbers correspond to the motifs detected in KA-1h. **K.** Signal for CREB, SRF and Fos binding 1 h after KCl stimulation of neuronal cultures (Malik et al., 2014) at the detected footprints 1 h after KA in IA promoters/enhancers. **L.** Conservation score for the detected footprints at IA promoter/enhancer regions.

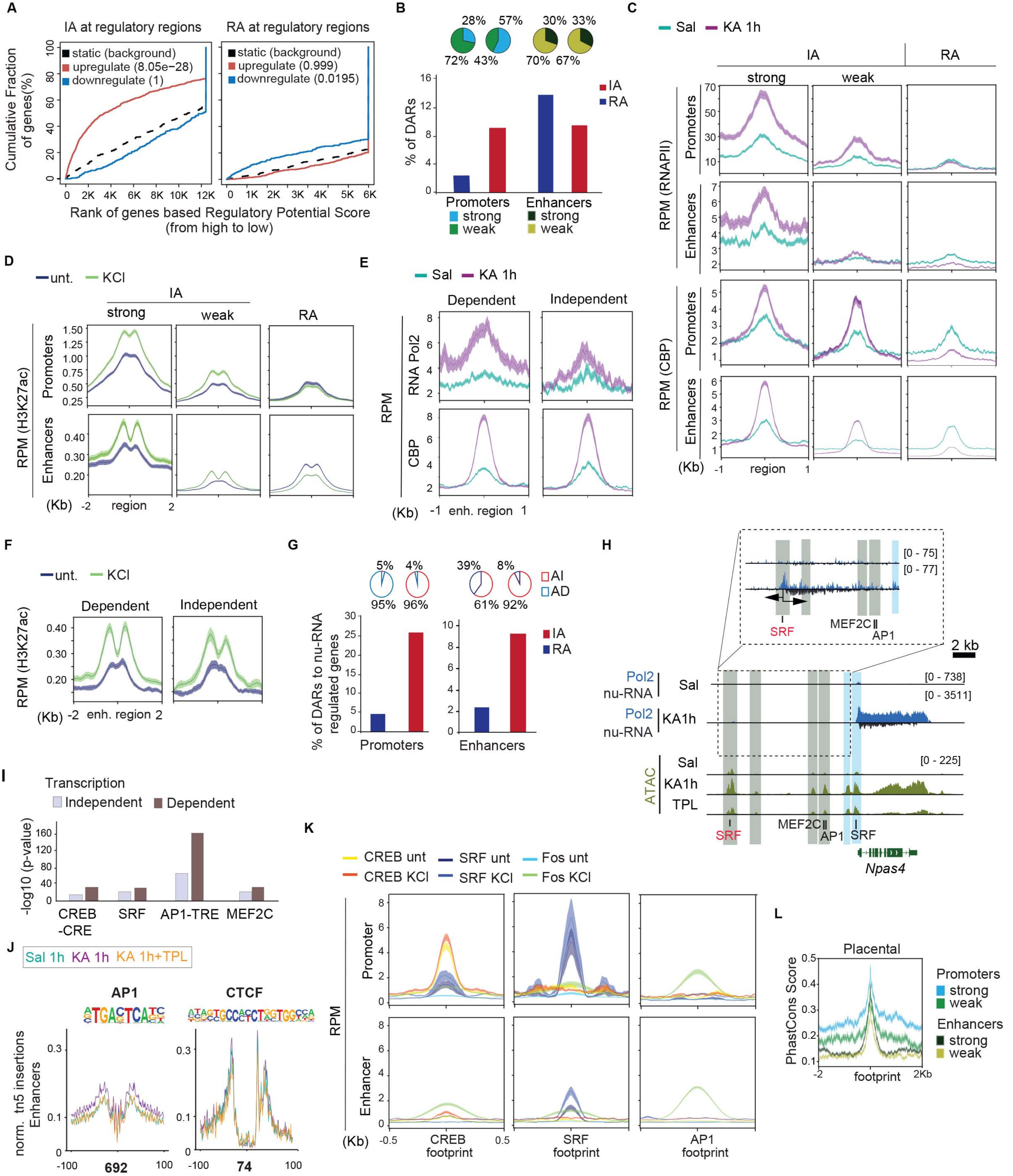

A
IA at regulatory regions
RA at regulatory regions
static (background)
upregulate (8.05e−28)
downregulate (1)
upregulate (0.999)
downregulate (0.0195)
Cumulative Fraction of genes(%)
Rank of genes based Regulatory Potential Score (from high to low)
B
28% 57% 30% 33%
72% 43% 70% 67%
% of DARs
IA
RA
Promoters
Enhancers
strong
weak
C
Sal
KA 1h
IA
RA
strong
weak
RPM (RNAPII)
RPM (CBP)
Promoters
Enhancers
(Kb) -1 region 1
D
unt.
KCl
RPM (H3K27ac)
(Kb) -2 region 2
E
Dependent
Independent
RNA Pol2
CBP
RPM
(Kb) -1 enh. region 1
F
(Kb) -2 enh. region 2
G
5% 4% 39% 8%
95% 96% 61% 92%
AI
AD
% of DARs to nu-RNA regulated genes
H
SRF
MEF2C
AP1
2 kb
Pol2 nu-RNA
Sal
KA1h
ATAC
TPL
[0 - 75]
[0 - 77]
[0 - 738]
[0 - 3511]
[0 - 225]
Npas4
I
Transcription
Independent
Dependent
-log10 (p-value)
CREB -CRE
SRF
AP1-TRE
MEF2C
J
Sal 1h
KA 1h
KA 1h+TPL
AP1
CTCF
norm. tn5 insertions Enhancers
692
74
K
CREB unt
CREB KCl
SRF unt
SRF KCl
Fos unt
Fos KCl
Promoter
Enhancer
RPM
(Kb) -0.5 0.5
CREB footprint
SRF footprint
AP1 footprint
L
Placental
PhastCons Score
Promoters
strong
weak
Enhancers
strong
weak
-2 footprint 2Kb

**Supp. Figure S5. Chromatin changes upon physiological neuronal activation.** **A**. Flow cytometry analysis of populations by its fluorescent intensities with the presence or absence of Fos (percentage of the filtered population indicated within the channel) after channel filtering following our protocol for the specific isolation of fluorescent singlets/nuclei in **Fig. S2A**. Right panel it is the co-localization of the size distribution of $Fos^+$ (purple) and $GFP^+$ cells (gray). **B.** Heat-map comparing DARs from SE and NE datasets, indicating common and exclusive regions between conditions**.** Last column is indicating the ratio of genic/intergenic regions on each subset. **C.** Principal component analysis for ATAC-seq datasets form physiologically ($Fos^+$ *vs* $Fos^-$ neurons) and chemically activated (Sal *vs* KA-1h) neurons. **D.** Volcano plot showing the significance value distribution after differential accessible regions analysis in $Fos^-$ neurons *vs* neurons from saline-treated mice. **E.** Predictive analysis of the accessibility changes at the regions annotated as promoter or enhancer and the change in expression of the gene (basic Activating/Repressive Function Prediction). AI: activity-induced genes; AD: activity-depleted genes; IA: increased accessibility regions; RA: reduced accessibility regions. **F.** Bar plot indicating the percentage of DARs in NE dataset at promoter and enhancer regions. **G.** Signal of activity regulated profiles 1 h after KA for RNAPII and CBP, comparing exclusive regions at detected enhancers' footprints in NE and SE. **H.** Signal for H3K4me1, H3K27ac and DNA methylation after fear conditioning (Halder et al., 2016) at detected NE-responding enhancers footprints.

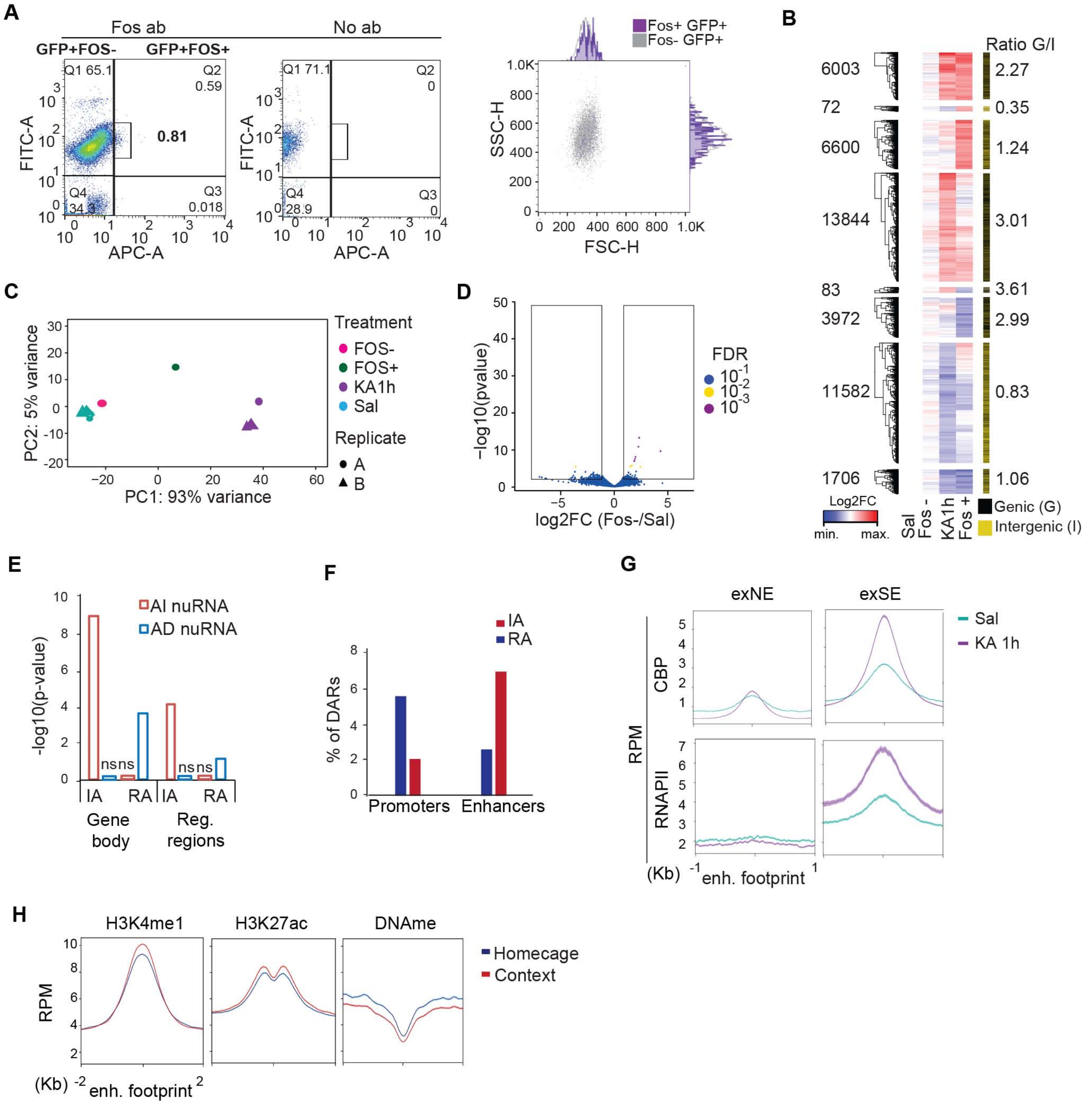

A
Fos ab
No ab
GFP+FOS-
GFP+FOS+
Q1 65.1
Q2 0.59
0.81
Q4 34.3
Q3 0.018
FITC-A
APC-A
Q1 71.1
Q2 0
Q4 28.9
Q3 0
Fos+ GFP+
Fos- GFP+
SSC-H
FSC-H
B
Ratio G/I
6003
2.27
72
0.35
6600
1.24
13844
3.01
83
3.61
3972
2.99
11582
0.83
1706
1.06
Log2FC
min.
max.
Sal
Fos -
KA1h
Fos +
Genic (G)
Intergenic (I)
C
Treatment
FOS-
FOS+
KA1h
Sal
Replicate
A
B
PC2: 5% variance
PC1: 93% variance
D
-log10(pvalue)
log2FC (Fos-/Sal)
FDR
E
AI nuRNA
AD nuRNA
-log10(p-value)
ns
IA
RA
Gene body
Reg. regions
F
IA
RA
% of DARs
Promoters
Enhancers
G
exNE
exSE
Sal
KA 1h
CBP
RNAPII
RPM
(Kb)
enh. footprint
H
H3K4me1
H3K27ac
DNAme
Homecage
Context
RPM
(Kb)
enh. footprint

**Supp. Figure S6. Hi-C analysis of activity-driven interactions. A.** Pearson correlation between Hi-C replicate samples at 0 h (saline), 1 h and 48 h after KA administration. **B.** Top: Snapshot of the *Fos* locus presenting *de novo* 4C promoter-enhancer interactions in response to SE. Red arrows indicate activity-driven interactions 1 h after KA. Bottom: Genomic profiles at the different time points. Values are in RPM. **C.** Metaplot of CTCF loops where at least one anchor overlaps a differential ATAC-seq peak indicating that CTCF loops do not change during neuronal activation despite changes to other transcription factors. IA: increased accessibility; RA: reduced accessibility. **D.** Venn Diagram of CTCF loops in KA-1h and saline samples. The low number of changes (< 4%, below the FDR level) indicates no real change. **E.** Top: Snapshot of the *Bdnf* locus presenting *de novo* 4C-interactions in response to SE. Red arrows indicate activity-driven interactions 1 h after KA. Bottom: Genomic profiles at the different time points. Values are in RPM. The right panel zooms in on the *Bdnf* gene body to show the increased interaction between the TSS and the TTS.

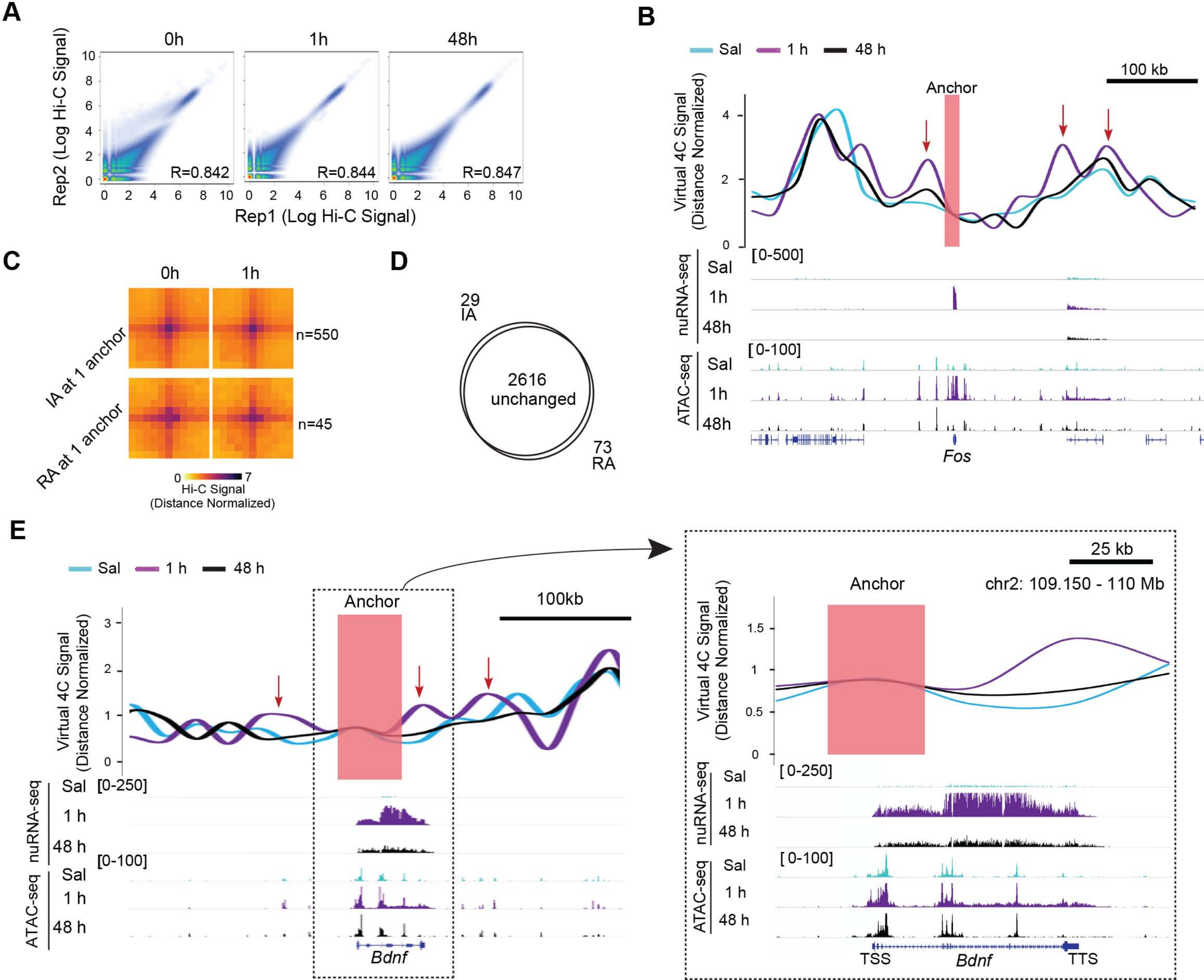
A
0h
1h
48h
Rep2 (Log Hi-C Signal)
Rep1 (Log Hi-C Signal)
R=0.842
R=0.844
R=0.847
B
Sal
1 h
48 h
100 kb
Anchor
Virtual 4C Signal (Distance Normalized)
nuRNA-seq
[0-500]
ATAC-seq
[0-100]
Fos
C
0h
1h
IA at 1 anchor
RA at 1 anchor
n=550
n=45
Hi-C Signal (Distance Normalized)
D
29 IA
2616 unchanged
73 RA
E
Sal
1 h
48 h
Anchor
100kb
Virtual 4C Signal (Distance Normalized)
nuRNA-seq
[0-250]
ATAC-seq
[0-100]
Bdnf
25 kb
chr2: 109.150 - 110 Mb
TSS
TTS

**Supp. Figure S7. Chromatin accessibility dynamics shows changes that stay long after the transcriptional burst. A.** Pearson correlation matrix between normalized samples clustered by Euclidian dendrogram for nuRNA-seq. **B.** Volcano plot showing the significance value distribution after differential gene expression analysis in nuRNA-seq samples after 0 h, 1 h, 6 h, 48 h from neuronal activation. AI: activity-induced; AD: activity-depleted. **C.** Venn diagram presenting the overlaps between the sets of DETs at each time point in the longitudinal nuRNA-seq analysis. Almost 20% of the changes observed 1 h after KA-treatment are still detected 5 h later, whereas only 2% remain 2 days later. **D.** Heatmap for FC in nuRNA-seq analysis. **E.** Heatmap of fold enrichment for biological process GO terms detected in AI and AD genes in the nucRNA-seq longitudinal analisis. The numbers indicate the number of genes associated with each term. **F.** Heatmap comparing DARs at the different time points**.** The last column (black/ yellow) shows the ratio of genic/intergenic regions on each subset. IA: increased accessibility; RA: reduced accessibility **G.** Pearson correlation matrix between normalized samples clustered by Euclidian dendrogram for ATAC-seq. **H.** Volcano plot showing the significance value distribution after differential accessible regions analysis in ATAC-seq samples after 0 h, 1 h, 6 h and 48 h of neuronal activation. IA: increased accessibility; RA: reduced accessibility. **I.** Venn diagram presenting the overlaps between the sets of DARs at each time point in the longitudinal ATAC-seq analysis.

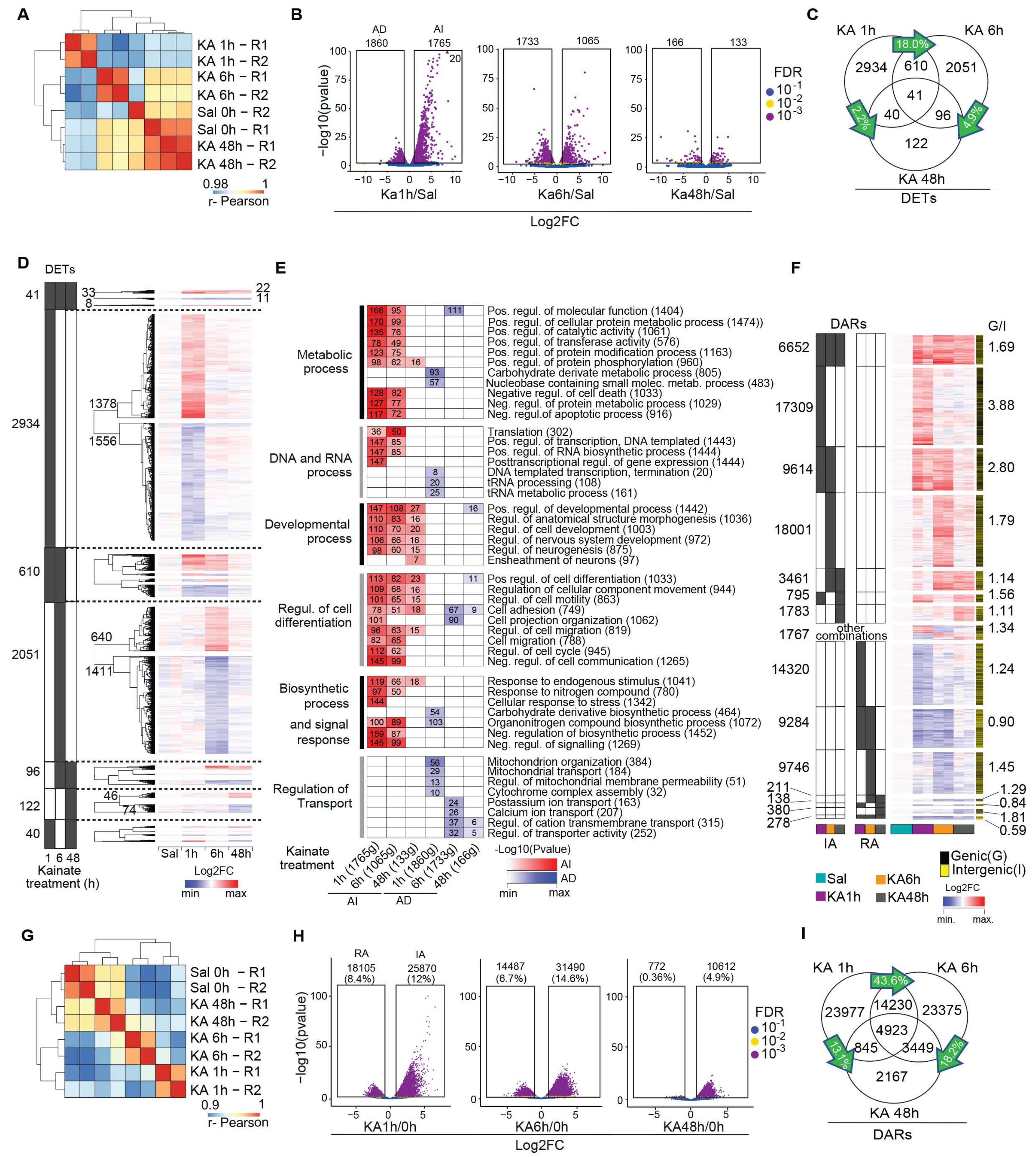

A
KA 1h – R1
KA 1h – R2
KA 6h – R1
KA 6h – R2
Sal 0h – R2
Sal 0h – R1
KA 48h – R1
KA 48h – R2
0.98
1
r- Pearson
B
AD
1860
AI
1765
20
1733
1065
166
133
−log10(pvalue)
Ka1h/Sal
Ka6h/Sal
Ka48h/Sal
Log2FC
FDR
$10^{-1}$
$10^{-2}$
$10^{-3}$
C
KA 1h
KA 6h
KA 48h
18.0%
2.2%
4.9%
2934
610
2051
41
40
96
122
DETs
D
DETs
41
2934
610
2051
96
122
40
33
8
22
11
1378
1556
640
1411
46
74
1 6 48
Kainate treatment (h)
Sal
1h
6h
48h
Log2FC
min
max
E
Metabolic process
Pos. regul. of molecular function (1404)
Pos. regul. of cellular protein metabolic process (1474))
Pos. regul. of catalytic activity (1061)
Pos. regul. of transferase activity (576)
Pos. regul. of protein modification process (1163)
Pos. regul. of protein phosphorylation (960)
Carbohydrate derivate metabolic process (805)
Nucleobase containing small molec. metab. process (483)
Negative regul. of cell death (1033)
Neg. regul. of protein metabolic process (1029)
Neg. regul.of apoptotic process (916)
DNA and RNA process
Translation (302)
Pos. regul. of transcription, DNA templated (1443)
Pos. regul. of RNA biosynthetic process (1444)
Posttranscriptional regul. of gene expression (1444)
DNA templated transcription, termination (20)
tRNA processing (108)
tRNA metabolic process (161)
Developmental process
Pos. regul. of developmental process (1442)
Regul. of anatomical structure morphogenesis (1036)
Regul. of cell development (1003)
Regul. of nervous system development (972)
Regul. of neurogenesis (875)
Ensheathment of neurons (97)
Regul. of cell differentiation
Pos regul. of cell differentiation (1033)
Regulation of cellular component movement (944)
Regul. of cell motility (863)
Cell adhesion (749)
Cell projection organization (1062)
Regul. of cell migration (819)
Cell migration (788)
Regul. of cell cycle (945)
Neg. regul. of cell communication (1265)
Biosynthetic process and signal response
Response to endogenous stimulus (1041)
Response to nitrogen compound (780)
Cellular response to stress (1342)
Carbohydrate derivative biosynthetic process (464)
Organonitrogen compound biosynthetic process (1072)
Neg. regulation of biosynthetic process (1452)
Neg. regul. of signalling (1269)
Regulation of Transport
Mitochondrion organization (384)
Mitochondrial transport (184)
Regul. of mitochondrial membrane permeability (51)
Cytochrome complex assembly (32)
Postassium ion transport (163)
Calcium ion transport (207)
Regul. of cation transmembrane transport (315)
Regul. of transporter activity (252)
Kainate treatment
1h (1765g)
6h (1065g)
48h (133g)
1h (1860g)
6h (1733g)
48h (166g)
AI
AD
-Log10(Pvalue)
AI
AD
min
max
F
DARs
G/I
6652
17309
9614
18001
3461
795
1783
1767
14320
9284
9746
211
138
380
278
1.69
3.88
2.80
1.79
1.14
1.56
1.11
1.34
1.24
0.90
1.45
1.29
0.84
1.81
0.59
other combinations
IA
RA
Genic(G)
Intergenic(I)
Sal
KA1h
KA6h
KA48h
Log2FC
min.
max.
G
Sal 0h – R1
Sal 0h – R2
KA 48h – R1
KA 48h – R2
KA 6h – R1
KA 6h – R2
KA 1h – R1
KA 1h – R2
0.9
1
r- Pearson
H
RA
IA
18105 (8.4%)
25870 (12%)
14487 (6.7%)
31490 (14.6%)
772 (0.36%)
10612 (4.9%)
−log10(pvalue)
KA1h/0h
KA6h/0h
KA48h/0h
Log2FC
FDR
$10^{-1}$
$10^{-2}$
$10^{-3}$
I
KA 1h
KA 6h
KA 48h
43.6%
13.1%
18.2%
23977
14230
23375
4923
845
3449
2167
DARs

**Supp. Figure S8. Long-lasting chromatin accessibility changes are associated with AP1 binding and disease. A.** Genomic screenshots for *Nptx2* (an example of activity-regulated gene that does not return to basal expression and maintains some proximal chromatin accessibility changes after 48 h) and *Klf4* (an example of activity-regulated gene that return to basal expression and maintains some proximal chromatin accessibility changes after 48 h). Footprint motifs for SRF, AP1, CRE, MEF2C are labeled. DArs are colored depending on their location in a promoter or enhancer. The maintained DARs after 48 h are labeled in green. In the case of *Klf4* we detect a non-annotated activity-regulated transcript produced from a bidirectional promoter with negative feedback regulation. Values show counts in RPM. **B.** Plot shows the footprinted sites at the maintained AP1 motifs detected in our longitudinal analysis (profile indicate the normalized tn5 insertions). **C.** Immunostaining for Fos in granular neurons at the DG and pyramidal neurons at the CA1 subfield at different time points after KA. **D.** Top: Scheme of object location memory (NOL) test. The time intervals from induced-recombination to SE and NOL testing are indicated. Bottom: Preference index for the displaced object in naïve mice and in mice that suffered SE one week earlier in the NOL test. **E.** RT-qPCR assay in RNA samples extracted 1 h after NE in animals that suffered SE two weeks earlier. FC values are referred to the home cage situation. **F.** Genomic screenshots for the *Fos/Jdp2 locus* (an example of activity-regulated gene that returns to basal expression but still maintains some proximal chromatin accessibility changes after 48 h). Labels are the same than in **panel A**.

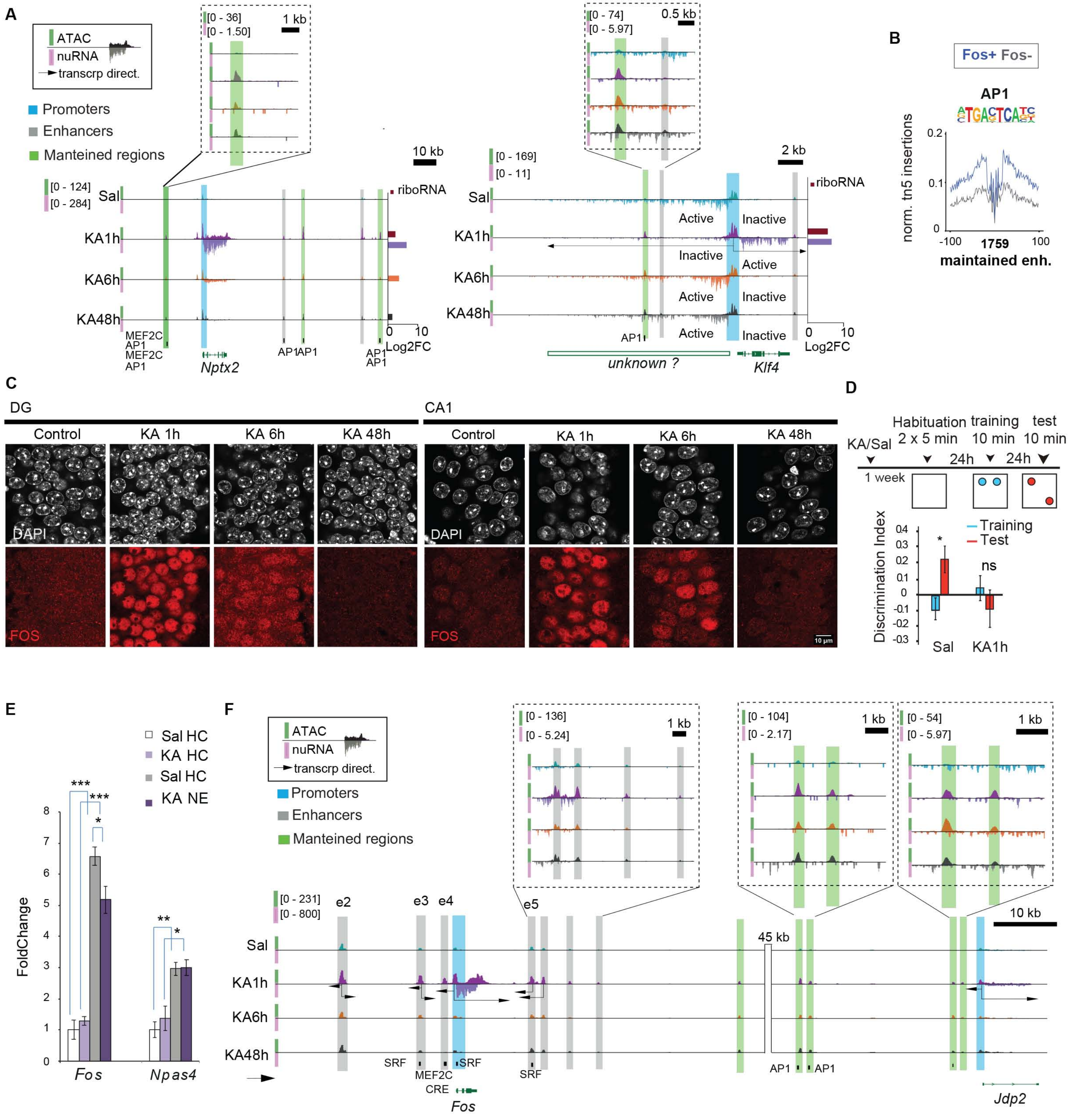

A
ATAC
nuRNA
transcrp direct.
Promoters
Enhancers
Manteined regions
[0 - 36]
[0 - 1.50]
1 kb
10 kb
[0 - 124]
[0 - 284]
Sal
KA1h
KA6h
KA48h
riboRNA
0 10
Log2FC
MEF2C
AP1
MEF2C
AP1
Nptx2
AP1 AP1
AP1
AP1
[0 - 74]
[0 - 5.97]
0.5 kb
[0 - 169]
[0 - 11]
2 kb
Active
Inactive
AP1
unknown ?
Klf4
B
Fos+ Fos-
AP1
norm. tn5 insertions
0.2
0.1
0
-100
1759
100
maintained enh.
C
DG
CA1
Control
KA 1h
KA 6h
KA 48h
DAPI
FOS
10 µm
D
Habituation
training
test
KA/Sal
2 x 5 min
10 min
10 min
24h
24h
1 week
Training
Test
Discrimination Index
*
ns
Sal
KA1h
E
Sal HC
KA HC
Sal HC
KA NE
***
***
*
**
*
FoldChange
Fos
Npas4
F
ATAC
nuRNA
transcrp direct.
Promoters
Enhancers
Manteined regions
[0 - 136]
[0 - 5.24]
1 kb
[0 - 104]
[0 - 2.17]
1 kb
[0 - 54]
[0 - 5.97]
1 kb
[0 - 231]
[0 - 800]
e2
e3
e4
e5
45 kb
10 kb
Sal
KA1h
KA6h
KA48h
SRF
MEF2C
SRF
CRE
SRF
AP1
AP1
Fos
Jdp2
